# Jasmonic Acid coordinates with Light, Glucose and Auxin signalling in Regulating Branching Angle of *Arabidopsis* Lateral Roots

**DOI:** 10.1101/2020.08.11.245720

**Authors:** Manvi Sharma, Mohan Sharma, K Muhammed Jamsheer, Ashverya Laxmi

**Author notes:** Corresponding author details: Ashverya Laxmi, Lab 203, National Institute of Plant Genome Research, Aruna Asaf Ali Marg, New Delhi-110067, India., 14, 17 Ext. 180. The authors contributed equally to this work (Manvi Sharma, Mohan Sharma).

## Abstract

The role of jasmonates (JAs) in primary root growth and development and in plant response to external stimuli is already known. However, its role in lateral root (LR) development remains to be explored. Our work identified methyl jasmonate (MeJA) as a key phytohormone in determining the branching angle of *Arabidopsis* LRs. MeJA inclines the LRs to a more vertical orientation, which was dependent on the canonical *JAR1-COI1-MYC2, 3, 4* signalling. Our work also highlights the dual roles of light in governing LR angle. Light signalling enhances JA biosynthesis, leading to erect root architecture; whereas, glucose (Glc) induces wider branching angles. Combining physiological and molecular assays, we revealed that Glc antagonizes the MeJA response via TARGET OF RAPAMYCIN (TOR) signalling. Moreover, physiological assays using auxin mutants, MYC2-mediated transcriptional activation of *LAZY2, LAZY4* and auxin biosynthetic gene *CYP79B2*, and asymmetric distribution of *DR5::GFP* and *PIN2::GFP* pinpointed the role of an intact auxin mechanism required by MeJA for vertical growth of LRs. We also demonstrated that light perception and signalling are indispensable for inducing vertical angles by MeJA. Thus, our investigation highlights antagonism between light and Glc signalling and how they interact with JA-auxin signals to optimize the branching angle of LRs.

## Introduction

The plant root system is highly plastic in nature and is influenced by exogenous and endogenous cues to modify its architecture. Because of their sedentary nature, plants are limited to their immediate surroundings for acquisition of water and nutrients (Malamy 2005; Cuesta, Wabnik & Benková 2013) and therefore, depend heavily on their root system. While the primary root maintains a nearly vertical orientation (orthogravitropism), lateral roots (LRs) form a non-vertical growth orientation, away from the main root axis, establishing a distinct gravitropic set-point angle (GSA) (Digby & Firn 1995). After their emergence, LRs adopt a perpendicular angle with respect to gravity, but as the LRs continues to develop, their GSA changes over time and exhibit de novo formation of an elongation zone (Kiss, Miller, Ogden & Roth 2002; Rosquete *et al*. 2013). The transient expression of PIN3 in LR columella cells defines asymmetric auxin distribution as well as differential elongation rates in stage II LRs (Rosquete *et al*. 2013), leading to the establishment of primary GSA of LR (Guyomarc’h *et al*. 2012) and is exemplified by an asymmetric growth towards gravity at a slower rate than the primary roots (Rosquete *et al*. 2013). As a result, the primary GSA of LRs is established. Subsequently, PIN3 repression in columella cells of stage III (SIII) LRs coincides with symmetric elongation, maintaining this primary GSA. Thus, the primary growth direction of LRs is of paramount significance, as it directs plant anchorage and uptake of water and nutrients, ultimately contributing to root system architecture.

In plants, phytohormones have profound effects on determining GSA. For example, as mentioned above, recent findings have identified the fundamental role of auxin in governing the GSA of shoot branches and LRs of *Arabidopsis thaliana* and other vascular plants, such as rice and beans (Rosquete *et al*. 2013; Rosquete, Waidmann & Kleine-Vehn 2018; Roychoudhry, Del Bianco, Kieffer & Kepinski 2013; Roychoudhry *et al*. 2017, 2019). Recently, Waidmann et al (2019) showed that CK signalling leads to reduced growth at the upper flank of the LR, thereby working as an antigravitropic component and hence preventing downward growth. In this way, a CK-dependent mechanism allows the root system to radially explore its surroundings (Waidmann *et al*. 2019). Jasmonic acid (JA) and methyl jasmonate (MeJA), collectively called jasmonates (JAs), are cyclopentenone compounds that are known to primarily modulate a number of vital physiological processes like root development, gravitropism, reproductive development, senescence; wound responses; and defence responses against pathogens and insects (Wasternack & Hause 2013). Findings of JA on regulation of main root gravitropism has already been reported (Gutjahr *et al*. 2005). It was found that the JA receptor CORONATINE-INSENSITIVE1 (COI1) is required for JA-mediated LR formation, positioning, and emergence of root bends in *Arabidopsis* (Raya-González, Pelagio-Flores & López-Bucio 2012). Additionally, investigations in rice have revealed the role of MeJA in modifying lamina joint inclination (Gan *et al*. 2015). Nonetheless, the role of JAs in steering LR GSA has not yet been demonstrated.

Besides phytohormones, factors such as temperature, nutrient status, and light play a vital role in governing branching angles (Digby & Firn 2002; Bai, Murali, Barber & Wolverton 2013; Trachsel, Kaeppler, Brown & Lynch 2013; Roychoudhry *et al*. 2017). Previous reports have implied the role of sucrose (Suc) in influencing the gravitropic behaviour of stems in *Paspalum vaginatum* (Willemoes, Beltrano & Montalbi 1988). Sugars such as Suc and glucose (Glc) are not only important nutrients, but also act as key signalling molecules (Ramon, Rolland & Sheen 2008; Eveland & Jackson 2012; Li & Sheen 2016). A recent report suggests that light and Suc act antagonistically to influence root length, but additively affect root hair emergence and elongation (García-González, Lacek & Retzer 2021). OsERF2 has been shown to influence rice root growth by fine tuning the genes involved in ABA and ethylene signalling pathways and sucrose metabolism (Xiao *et al*. 2016). Sucrose also modulates auxin metabolism, transport and signalling to regulate elongation growth responses (Lilley, Gee, Sairanen, Ljung & Nemhauser 2012; Sairanen *et al*. 2013; Stokes, Chattopadhyay, Wilkins, Nambara & Campbell 2013; Lastdrager, Hanson & Smeekens 2014). In *Arabidopsis*, there are three distinct Glc signalling pathways: (1) the HEXOKINASE1 (HXK1)-dependent pathway that is governed by HXK1-mediated signalling function, in which Glc is possibly sensed by a module independent of the catalytic activity of HXK1; (2) the G-protein-coupled receptor signalling by REGULATOR OF G-PROTEIN SIGNALLING 1 (RGS1) and GPA1 which are implicated in sensing extracellular glucose and signalling through THF1, located in the plastids (Huang *et al*. 2006; Urano *et al*. 2012a); and (3) a glycolysis-dependent pathway that works through the antagonistic interaction between SUCROSE NONFERMENTING RELATED KINASE 1 (SnRK1) and TARGET OF RAPAMYCIN (TOR) (Baena-González, Rolland, Thevelein & Sheen 2007; Baena-González 2010; Xiong & Sheen 2015; Song *et al*. 2017). Previous studies have suggested the involvement of Glc in various aspects of early seedling development (Mishra, Singh, Aggrawal & Laxmi 2009; Kircher & Schopfer 2012; Yuan, Xu, Zhang, Guo & Lu 2014). Glc and phytohormones have been extensively shown to interact with each other to enhance plant fitness. The interplay of Glc with various phytohormones has been shown to modulate root directional responses in *Arabidopsis* seedlings (Singh, Gupta & Laxmi 2014a b). However, very few reports link JA and sugar signalling (Song *et al*. 2017; Vleesschauwer *et al*. 2017; Guo *et al*. 2018). Based on physiological and pharmacological studies, an intimate cross-talk occurs between TOR and JA signalling pathways at multiple levels of JA signal transduction (Vleesschauwer *et al*. 2017), but the JA-Glc signal crosstalk that regulates branching angle of *Arabidopsis* LRs still remain obscure.

Despite the obvious importance of directional LR growth for plant propagation; the role of factors other than CK and auxin are largely unexplored. In this study, we have characterized the interaction of JA and Glc in the regulation of branching angle of LRs in *Arabidopsis*. Using genetic, physiological, molecular and cell biological approaches, we show that two opposing signals (JA and Glc) counteract each other in order to set directional LR growth. Additionally, knowledge gained from this study can expand our insight on root architecture establishment and may contribute to the regulation of GSA in the field of plant breeding.

## Materials and methods

### Plant materials

*Arabidopsis thaliana* ecotype*s* of Col-0, Ws and Ler were used as wild-type controls. The following seed stocks were obtained from the *Arabidopsis* Biological Resource Center (ABRC) at Ohio State University (http://www.Arabidopsis.org/abrc/): DR5::GFP; *phyA-201* (CS6291); *phyB-5* (CS6213); *phyA201B5* (CS6224); *hy5-1* (CS71); *rgs1-1* (CS6537); *rgs1-2* (CS6538); *gpa1-3* (CS6533); *gpa1-4* (CS6534); *tir1-1* (CS3798); *axr1-3* (CS3075); *aux1-7* (CS3074); *axr2-1* (CS3077); *eir1-1* (CS8058); *myc2/jin1-9* (SALK_017005C); *coi1* (SALK_095916C); *jar1-11* (CS67935); *jaz1* (SALK_011957C); *jaz2* (SALK_012782C); *jaz4* (SALK_141628C); *jaz6* (SALK_017531); *sweet11sweet12* (CS68845). Following lines were obtained from the original published source as: *myc2myc3myc4* (Schweizer et al., 2013); *mdr1-1* (Noh et al., 2001); *lax3* (Swarup et al., 2008); *pin3-4, pin4-3* (Friml et al., 2002b); *pin7-2* (Friml et al., 2003); *tor35-7 RNAi* (Deprost *et al*. 2007); *p35S::Jas9-N7-VENUS* (Larrieu et al., 2015); *35S::MYC2::GFP* (Jung et al., 2105); *PIN2::PIN2-eGFP* (NASC); *PIN3::PIN3-eGFP* (NASC). All mutant lines were in the Col-0 background except the following: *mdr1-1* was derived from Ws background; and *hy5-1, phyA-201, phyB-5, phyA201B5* were in the Ler background.

### Growth conditions

Seeds were surface sterilized and stratified at 4°C for 48 hours. The imbibed seeds were grown vertically on square petri dishes containing 1/2 Murashige and Skoog (MS) medium supplemented with 1% sucrose (29.13 mM; w/v) and solidified with 0.8% agar (w/v). Seed germination and plant growth were carried out in climate-controlled growth rooms under long day conditions (16 hr light and 8 hr darkness), with 22°C ± 2°C temperature and 60 μmol/sec/m^2^ light intensity. To study branching angle of LRs, five-day-old MS grown seedlings were transferred to hormone/inhibitor/sugar treatment media with their root tips marked and grown vertically under above mentioned growth conditions, unless otherwise stated. All chemicals were purchased from Sigma (St. Louis, MO, USA) except specified otherwise. MeJA was prepared as 50 mM stock solution in 100% (v/v) ethanol. Epibrassinolide was prepared as 10^−2^ M stock solution in 50% (v/v) ethanol. The following were prepared as 10^−2^ M stock solutions in dimethyl sulfoxide: NPA, BAP, ABA, and GA3. ACC was prepared as a sterile 10^−2^ M aqueous stock solution.

### Physiological analyses

Seedlings were grown on 1/2 MS, 0.8% agar, and 1%Suc-containing medium for five days. They were then transferred to 1/2 MS medium, supplemented with different combinations of Glc [0.5% (27.75 mM; w/v) and 3% (166.52 mM; w/v)] and MeJA (10 μM) or 1/2 MS medium supplemented with 1% sucrose and MeJA (1 μM, 5 μM, 10 μM) or combinations of Glc (1%) and Man (1%, 2%, 3%, 4%) with and without 10 μM MeJA and their root tips were marked. Dark experiments were carried out by transferring light grown, five-day-old WT and *myc2myc3myc4* seedlings to treatment media in light for 24 hours, followed by complete darkness for the next 5 days. For measurement of branching angle, the angle formed between each LR and primary root was measured. The average branching angle was determined by summing the angle formed by each LR of all the seedlings divided by the total number of LRs. Also, the angular response was measured by distributing the branching angle in three categories viz. <40°, 40°-70°, >70° and expressed as percentage of LR. ImageJ program from NIH was used to quantify the branching angle as well epidermal cell length of LRs (http://rsb.info.nih.gov/ij/). Digital images were captured from Nikon Cool pix camera on 12^th^ day of seedling growth.

### Laser Confocal Scanning Microscopy

To observe auxin distribution, PIN2 and PIN3 expression in stage II LRs, five-day-old light grown *DR5::GFP, PIN2::PIN2-eGFP* and *PIN3::PIN3-eGFP* expressing seedlings were transferred to 1/2 MS and 1/2 MS + 10 μM MeJA for 6-7 days for imaging. GFP fluorescence was imaged under a Leica TCS SP8 AOBS Laser Confocal Scanning Microscope (Leica Microsystems). To image GFP, the 488 nm line of the argon laser was used for excitation and emission was detected at 520 nm. The laser, pinhole and gain settings of the confocal microscope were kept identical among different treatments. Images were assembled using Photoshop (Adobe Systems). For optical sectioning under SP8 microscope, Z-stacks were scanned for 6 sections every 4 μm in thickness and maximum projections were generated. At least two biological replicates, with each replicate having 15 seedlings, were performed for all the experiments.

### Gene Expression Analysis

For gene expression analysis, RT-qPCR was performed. Imbibed seeds of WT, *myc2myc3myc4*, and *hy5-1* were sown on 1/2 MS medium supplemented with 1% (w/v) Suc and 0.8% (w/v) agar and grown vertically in culture room conditions. Five-day-old light grown seedlings of WT and *myc2myc3myc4* were harvested and stored in -80°C. WT and *hy5-1* were grown in light (long day regime) and continuous dark for 5 days. For dark to light transition, WT and *hy5-1* were grown in continuous dark for 5 days followed by light treatment for 6 hours. To check, JA related gene expression, 5-d–old Col-0 seedlings were first starved under 0% Glc for 24 h in the dark. After starvation, seedlings were treated without or with Glc and a combination of 10 μM MeJA. Afterwards, whole seedlings were flash frozen in liquid nitrogen and stored at -80°C. Total RNA was isolated from frozen tissue using the RNeasy Plant Mini Kit (Qiagen) following the manufacturer’s protocol. RNA was quantified and tested for quality before it was used for subsequent analyses. First-strand cDNA was synthesized by reverse transcription using 2 μg of total RNA with a high-capacity cDNA Reverse Transcription Kit (Applied Biosystems). Primers were designed by identifying a sequence stretch with a unique sequence. All candidate gene primers were designed using the software Primer Express (version 3.0; Applied Biosystems). An ABI Prism 7900 HP fast real-time PCR system (Applied Biosystems) was used. For the normalization of variance among samples, UBIQUITIN10 (UBQ10) was used as a reference control. The fold-change for each candidate gene in different experimental conditions was determined using the quantitative ΔΔCT method. All primers used are mentioned in the Supplemental Table S1. Gene expression analysis was performed 3 times unless otherwise stated.

### Chromatin Immunoprecipitation

ChIP assays were performed by following the protocol of Saleh et al. (2008) with minor modifications. *Arabidopsis* Col-0 and *35S::MYC2::GFP* seedlings were grown for seven days in 1/2 MS containing 1%Suc and used for the ChIP assay. Briefly, 1 gram (g) tissue of each sample was cross-linked with 1% formaldehyde to fix protein-DNA complexes. The samples was crushed in liquid N2 and homogenized in nuclei isolation and nuclei lysis buffers followed by sonication. Sonication of chromatin was done in 4°C water sonicator (Diagenode Bioruptor Plus). Sonicated samples were first precleared with Protein A Agarose beads (Millipore #16-157) and then mixed with antibodies. Antibodies against GFP (Cat. no. ab290) were purchased from Abcam. Approximately 200 bp upstream promoter region of *LAZY2, LAZY4* and *CYP79B2* containing the G- and E-box elements which are known binding sites of MYC2 was enriched in the ChIP assay. All primers used in the ChIP assays are mentioned in the Supplementary Table S1.

### Protein extraction and immunoblot assay

Extraction of soluble proteins was performed on seven day old seedlings of p35S::Jas9-VENUS. Seven day old light grown seedlings of p35S::Jas9-N7-VENUS were treated in sugar-free liquid 1/2 MS in dark for 24 h to deplete internal sugars. This was followed by treating the seedlings with 0% (w/v) Glc and 3% (w/v) Glc containing 1/2 MS liquid medium for 3 hours. Following Glc treatment, 10 μM MeJA or 10 μM AZD-8055 or both were added to all seedlings for 3 hours in dark. Seedlings were harvested, frozen in liquid nitrogen and then ground in pre chilled mortar pestle using liquid nitrogen. The powder was resuspended in cold extraction buffer (137 mM of NaCl, 2.7 mM of KCl, 4.3 mM of Na2HPO4, 1.47 mM of KH2PO4, 10% glycerol [1.1 M], and 1 mM phenylmethylsulfonyl fluoride [PMSF]) supplemented with plant protease inhibitor cocktail (Sigma-Aldrich, http://www.sigmaaldrich.com/). This was followed by two rounds of centrifugation (15□min, 13,000□rpm at 4□°C) to remove cell debris. The samples were boiled for 10□min at 95□°C before being loaded onto a gel (40□μl; 80-100 μg per lane). SDS–PAGE was performed on 10% polyacrylamide gel. After transfer onto a PVDF membrane, protein amounts in each lane were checked using Ponceau staining (0.1%, 1.5 mM). Immunoblots were detected using a primary rabbit polyclonal anti-GFP antibody (ab290, Abcam, diluted 1:5,000), primary rabbit polyclonal anti HSP90-2 antibody (diluted 1:5,000) and a secondary anti-rabbit IgG-HRP (diluted 1:10,000). Proteins were visualized using the enhanced chemiluminescence kit.

### Microarray analysis

For microarray of *Arabidopsis* WT seedlings under Glc and MeJA treatment, 5-d–old seedlings were first starved under 0% Glc for 24 h in the dark. After starvation, seedlings were treated with and without Glc and a combination of 10 μM MeJA. Harvested samples were outsourced for transcriptome profiling. The data were analyzed by the Transcriptome Analysis Console (v3.0; Affymetrix) with default parameters. Two biological replicates were used for the microarray experiment.

### Statistical analyses

All physiological experiments yielding similar results were repeated as mentioned in the figure legends, in which each experiment was considered as an independent biological replicate consisting of at least 25 seedlings. Immunoblot assays were performed 4 times unless otherwise stated. ChIP assays were performed as mentioned in the figure legends. Statistical differences between control/treatment and WT/mutant pair were analyzed using Student’s *t* test with paired two-tailed distribution. *P*-value cutoff was taken at *P* <0.05. All data were managed and analyzed using Microsoft Excel. The graphs were made using Microsoft Excel and Instant Clue. End point physiological analyses were carried out on 12^th^ day of seedling growth.

### Accession Numbers

*Arabidopsis* Genome Initiative locus identifiers for the genes mentioned in this article are: *phyA-201*, AT1G09570; *phyB-5*, AT2G18790; *hy5-1*, AT5G11260; *tor35-7 RNAi* AT1G50030; *rgs1-1*, AT3G26090; *rgs1-2* AT3G26090; *gpa1-3* AT2G26300; *gpa1-4*, AT2G26300; *tir1-1*, AT3G62980; *axr1-3*, AT1G05180; *aux1-7*, AT2G38120; *axr2-1*, AT3G23050; *eir1-1*, AT5G57090; *myc2/jin1-9*, AT1G32640; *coi1*, AT2G39940; *jar1-11*, AT2G46370; *jaz1*, AT1G19180; *jaz2*, AT1G74950; *jaz4*, AT1G48500; *jaz6*, AT1G72450; *mdr1-1*, AT3G28860; *lax3*, AT1G77690; *pin4-3*, AT2G01420; *pin7-2*, AT1G23080; *pin3-4*, AT1G70940; *sweet11swet12*, (AT3G48740, AT5G23660).

## Results

### MeJA modulates branching angle of *Arabidopsis* roots

To study the role of MeJA in regulating growth angles, the branching angle of *Arabidopsis* WT LRs was analysed by two methods: 1) calculating the angle formed by all LRs of wild-type (WT) seedlings with respect to the main root and averaging the values, and 2) distributing the angles of LRs in three categories (<40°, 40°-70° and >70°). We observed that MeJA treatment decreases the branching angle in a dose-dependent manner (Fig. 1A-C). LRs of WT showed an average branching angle of 67.02° on control media, but a more vertical angle of 40.21° when grown on the high MeJA concentration (10 μM) (Fig. 1B). Approximately 63% of LRs adopted angles between 0-40° and very few with angles >70° when grown on 1/2 MS supplemented with 10 μM MeJA (Fig. 1C). Since the effect of auxin and CK in governing GSA is already explored (Rosquete *et al*. 2013, 2018; Roychoudhry *et al*. 2013, 2019; Waidmann *et al*. 2019), we wanted to identify the role of other hormones in modulating this response. The *Arabidopsis* WT seedlings did not display any changes in branching angle when grown in brassinosteroids (BR), gibberellic acid (GA_3_), 1-aminocyclopropane-1-carboxylic acid (ACC) or abscisic acid (ABA) (Fig. 1D). Altogether, our data indicates that MeJA impacts the angular growth of LRs.

**Figure 1.**
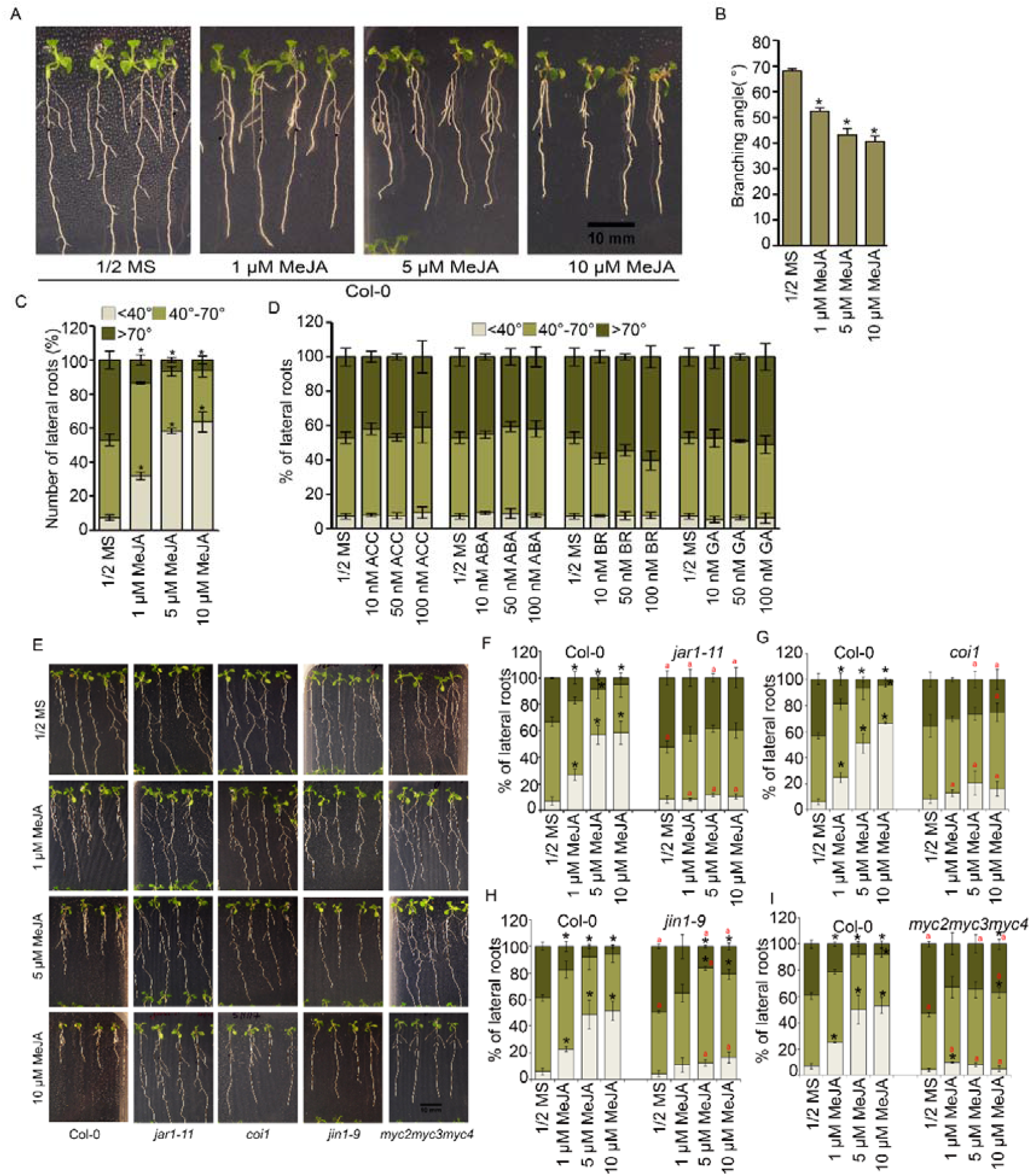
MeJA decreases branching angle of Arabidopsis roots. **(A)** Phenotype of light grown 12-day-old Arabidopsis Col-0 seedlings grown on different doses of MeJA. **(B)** Average branching angle of 12-day-old Col-0 seedlings on different doses of MeJA. Data are mean-SE of 3 biological replicates with 25 seedlings. **(C)** Distribution of branching angle of 12-day-old Col-0 Arabidopsis seedlings on different doses of MeJA. Data are mean-SE of 3 biological replicates with 25 seedlings. **(D)** Comparison of the effects of different phytohormones (ACC, ABA, BR, GA) on Col-0 seedlings to determine their roles in branching angle. Data are mean-SE of 3 biological replicates with 25 seedlings. **(E)** Phenotype of light grown 12-day-old Arabidopsis Col-0, *jar1-11, coi1, jin1-9* and *myc2myc3myc4* seedlings grown on different doses of MeJA. **(F)** Distribution of branching angle in Col-0, *jar1-11* seedlings grown on different doses of MeJA. Data are mean-SE of 3 biological replicates with 25 seedlings. **(G)** Distribution of branching angle in seedlings of Col-0, *coi1* mutants grown on different doses of MeJA. Data are mean-SE of 4 biological replicates with 25 seedlings. **(H-I)** Distribution of branching angle in seedlings of Col-0, *jin1-9, myc2myc3myc4* mutants grown on different doses of MeJA. Data are mean-SE of 3 biological replicates with 25 seedlings. In A-I, 5-day-old 1/2 MS grown Col-0 seedlings were transferred to different doses of MeJA and phenotypes were analysed at 12^th^ day. Asterisks indicate a significant difference in the studied parameter (P <0.05, paired two-tailed student’s t-test; * control vs treatment and ‘a’ WT vs mutant).

### MeJA decreases branching angle in a SCF^COI1^ and MYC2, 3, 4 dependent pathway

In order to elucidate how JA biosynthesis and signalling are involved, several mutants defective in the JA-biosynthesis and signalling pathways were assessed. JA perception by COI1 is the first committed step of JA signalling. Before perception, JA is converted to its biologically active form, jasmonoyl-isoleucine (JA-Ile), by the enzyme JASMONATE RESISTANT 1 (JAR1) (Staswick & Tiryaki 2004; Staswick 2009). Neither *jar1-11* nor *coi1* responded to MeJA treatment, and both exhibited an overall horizontal orientation of LRs (Fig. 1E-G). Also, very few LRs of these mutants adopted angles <40°, as compared to WT (Fig. 1E-G). The average branching angles of *jar1-11* and *coi1* indicated a complete resistance to the MeJA treatment (Fig. S1A). MYC2 is a bHLH transcription factor that is considered the master regulator of jasmonate and light responses (Lorenzo, Chico, Sá Nchez-Serrano & Solano; Dombrecht *et al*. 2007). Therefore, we assessed whether JA-dependent transcription factors indeed have an impact on LR growth in the WT. The *jasmonate insensitive 1* (*jin1-9)*, a mutant of *MYC2*, was less responsive to MeJA treatment as compared to WT, as its LR showed broader angles (Fig. 1E, H; Fig. S1B). Also, *myc2myc3myc4* displayed diminished sensitivity, as a large percentage of LRs acquired angles >70*°* as compared to the single mutant and WT (Fig. 1E-I; Fig. S1B), suggesting that *MYC3* and *MYC4* act additively with *MYC2* in regulating this response. T-DNA insertion lines of *JASMONATE-ZIM-DOMAIN* (*JAZ1, JAZ2, JAZ4*, and *JAZ6*) did not show any major changes to increasing MeJA doses which can be attributed to redundancy of *JAZ* genes (Fig. S1C).

Since our in vitro study allowed only two-dimensional analysis of root growth, we further assessed angular growth of LRs in three dimensional systems. To allow three dimensional root expansion in vitro, we cultivated twenty-days-old WT, *jar1-11* and *myc2myc3myc4* plants in cylindrical tubes containing 1/2 MS with 1% phytagel, and found broader LR angles when compared to WT (Fig. S2A and B), suggesting that our two-dimensional in vitro screen was highly suitable for identifying mutants with diverging GSA values. Moreover, these results suggest that the defects in endogenous JA biosynthesis and signalling machinery leads to broader LR angle. Altogether, this set of data confirms that JA signalling utilizing receptor COI1 and TF MYC2 regulates angular growth of LRs.

### Glc production via photosynthesis negatively influences MeJA-modulated branching angle

Previously, Digby and Firn (2002) demonstrated that light effects on GSA of organs can be brought about both via the action of photosynthesis and via phytochrome reception and signalling (Digby & Firn 2002). Previous reports have established the role of Glc in primary root gravitropism in *Arabidopsis* (Singh *et al*. 2014a, b). In order to explore the role of Glc and its interaction with MeJA in altering branching angles, the WT seedlings were treated with MeJA (10 μM) in combination with different concentrations of Glc (0.5%Glc; 28 mM and 3%Glc; 167 mM). The 3% concentration of Glc widened the branching angle as compared to the 0.5% concentration (Fig. 2A-B; Fig. S3). The LRs attained more vertical orientation when treated with a combination of 0.5%Glc+MeJA. However, when both MeJA and 3%Glc were added together, MeJA still showed a strong effect of narrowing root angle in the presence of 3% Glc, but the overall percentage of root angles did not reach the same level as they do in the 0.5% treatment (Fig. 2A-B; Fig. S3A). The above results suggest an antagonistic interaction between Glc and MeJA in controlling this response. We also used 5%Glc to check whether higher Glc concentration causes more widening of branching angle. However, we did not observe any significant changes in the LR angle between 0.5%Glc and 5%Glc, thereby suggesting that 3%Glc is the optimal concentration for plant growth that widens LR angle (Fig. S3B-C). Also, higher Glc concentrations is shown to cause developmental arrest in WT seedlings such as anthocyanin accumulation, less LR production and shorter root length (Moore *et al*. 2003) and therefore is considered inhibitory for overall growth and development of plants. Next, we wanted to assess whether the observed phenotype is due to changes in the osmoticum of the growth media or specific to Glc. To test this, we used mannitol (Man), a non-metabolizable sugar-alcohol that initiates very few LRs (Gupta, Singh & Laxmi 2015). Hence, to induce LR production, a small amount of Glc (1%) was also combined with increasing concentrations of Man (1%, 2%, 3% and 4%). We observed that the effects previously induced by the supply of Glc were not produced by Man. Increased doses of Man induced a vertically oriented LRs as compared to Glc (Fig. 2C). Thus, osmotic changes do have an effect on branch angle, however it appears to be the opposite effect than that of Glc. SUGARS WILL EVENTUALLY BE EXPORTED TRANSPORTERS (SWEET) are bidirectional sugar uniporters that are capable of transporting Suc, Glc and Fruc (Chen *et al*. 2010). In order to test whether changes in endogenous sugar level in root alters the LR angle, we used the *sweet11sweet12* mutants defective in sugar transport from shoot to root. For this, we transferred 5-day-old 1/2 MS grown seedlings on low (0.5%) and moderate (1%) Glc concentration and observed that *sweet11sweet12* showed steeper branching angles as compared to WT in both Glc concentrations (Fig. 2D-E). This result indicates that depletion of endogenous Glc in root alters the LR branching angle suggesting the role of Glc in inducing wider branching angle in *Arabidopsis*.

**Figure 2.**
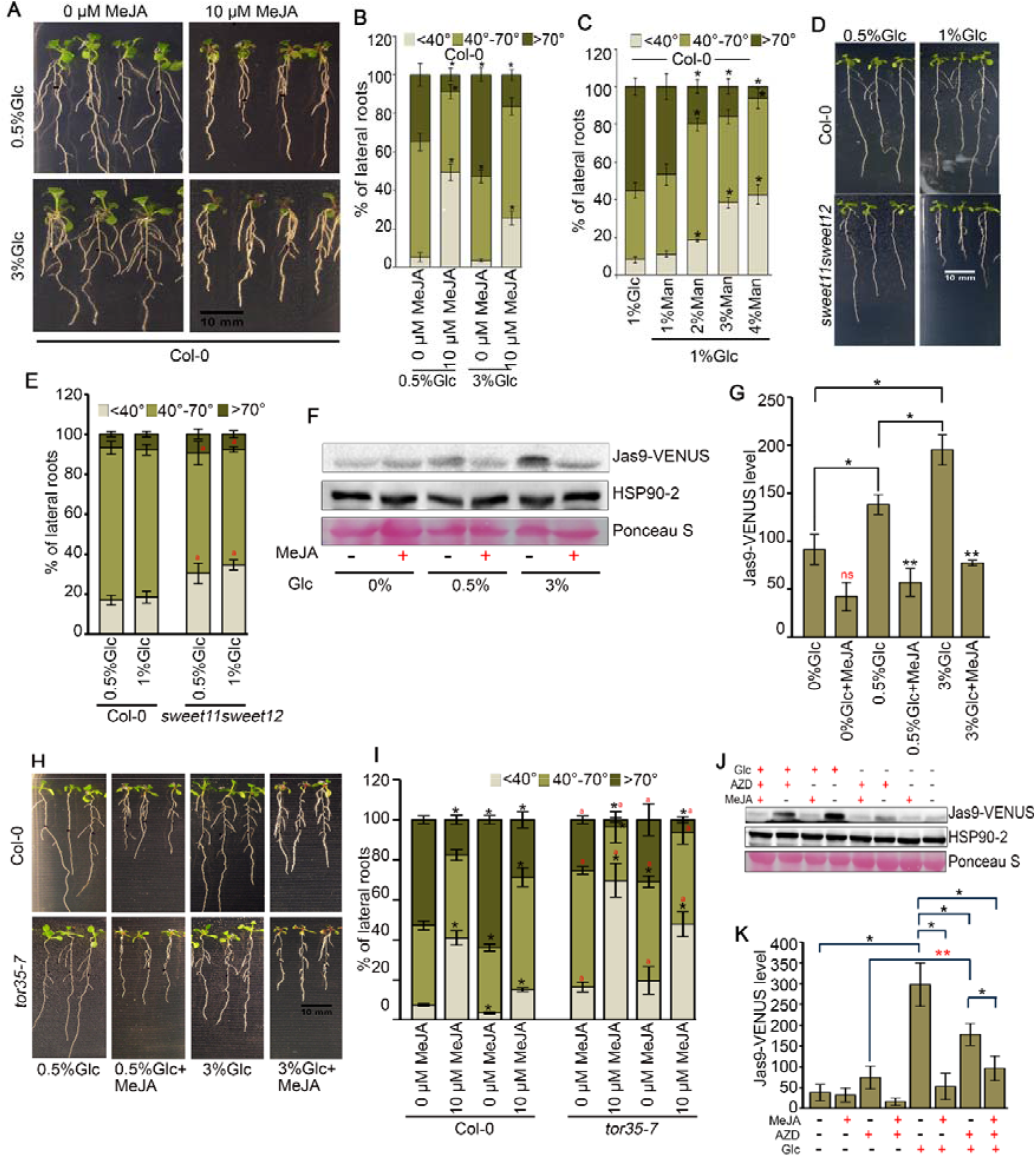
Glc antagonizes MeJA-induced branching angle in Arabidopsis roots. **(A)** Phenotype of Col-0 seedlings grown in different concentrations of Glc (0.5%; 28 mM and 3%Glc; 167 mM) and in combination with 10 μM MeJA. **(B)** Distribution of branching angle in Col-0 seedlings grown in different concentrations of Glc and in combination with MeJA. The graph represents the average of 7 biological replicates having 25 plants in each replicate and error bars represent SE. **(C)** Distribution of branching angle in Col-0 seedlings grown in different concentrations of Man in combination with 1%Glc. The graph represents the average of 3 biological replicates consisting of 25 seedlings and error bars represent SE. **(D)** Phenotype of Col-0 and *sweet11sweet12* seedlings grown in different concentrations of Glc. **(E)** Distribution of branching angle in Col-0 and *sweet11sweet12* seedlings grown in different concentrations of Glc. The graph represents the average of 3 biological replicates having 50 plants in each replicate and error bars represent SE. In **A-E**, 5-day-old 1/2 MS grown seedlings were transferred to Glc and MeJA containing treatment media and phenotypes were analysed at 12^th^ day. Asterisks indicate a significant difference in the studied parameter (*P* <0.05, Student’s t-test; * control vs treatment and ‘a’ WT vs mutant. **(F)** Western blot analysis of total protein extracts of Jas9-VENUS seedlings treated for 3 hours with 0%, 0.5% and 3%Glc in combination with 10 μM MeJA and probed with an anti-GFP antibody. The experiment was performed four times. **(G)** Quantification of Jas9-VENUS protein levels. The graph represents average of 4 biological replicates. **(H)** Phenotype of Col-0 and *TOR* signalling mutant *tor35-7 RNAi* seedlings grown in different concentrations of Glc alone and in combination with 10 μM MeJA. **(I)** Distribution of branching angle in seedlings of Col-0 and *tor35-7 RNAi* grown in different concentrations of Glc alone or in combination with 10 μM MeJA. 5-day-old 1/2 MS grown Col-0 and *tor35-7* seedlings were transferred to treatment media and phenotypes were analysed at 12^th^ day. The graph represents the average of 4 biological replicates consisting of 25 seedlings and error bars represent SE. Asterisks indicate a significant difference in the studied parameter (*P* <0.05, Student’s t-test; * control vs treatment, and ‘a’ WT vs mutant). **(J)** Western blot analysis of total protein extracts of Jas9-VENUS seedlings treated for 3 hours with 0%, 0.5% and 3%Glc alone or in combination with 10 μM MeJA and 10 μM AZD-8055 for 3h and probed with an anti-GFP antibody. The experiment was performed four times. **(K)** Quantification of Jas9-VENUS protein levels. The graph represents average of 4 biological replicates. Asterisks indicate a significant difference in the studied parameter (P <0.05, paired two-tailed student’s t-test; * control vs treatment and ** 0%Glc+AZD vs 3%Glc+AZD.

Next, to understand the crosstalk between Glc and JA signalling in influencing LR angle at the molecular level, we first examined the protein level of JAZ9, a negative regulator of JA signalling, in the presence and absence of Glc. For this, we used Jas9-VENUS, a fluorescent marker widely used for the perception of bioactive JA, showing dynamic changes in JA levels (Larrieu *et al*. 2015). As shown in Figure 2F-G, we observed very low levels of Jas9-VENUS protein in seedlings treated with 0%Glc and 0%Glc+MeJA. Interestingly, as the concentration of Glc was increased from 0 to 0.5% and 3%, Jas9-VENUS fusion protein was accumulated at higher levels (Fig. 2F-G; Fig. S4A), which was found to be degraded when subjected to MeJA (3%Glc+MeJA). We also found that even a low amount of Glc was sufficient for MeJA mediated degradation of Jas9-VENUS (Fig. 2F-G; Fig. S4A). To test whether MeJA can be perceived properly in the absence of sugar in the media, we compared the expression of various JA-related genes (*JAZ1, JAZ2, JAZ3, JAZ6, JAZ9, AOS, ERF1, ORA59* and *MYC2*) in WT seedlings subjected to MeJA treatment without Glc using RT-qPCR and found out that these genes were upregulated in MeJA treatment even without Glc (Fig. S4B). This suggests that MeJA is actively perceived within the system even in the absence of sugar. (Fig. 2F-G; Fig. S4A). Thus, all these results suggest that Glc promotes the accumulation as well as MeJA-mediated degradation of Jas9-VENUS protein.

### Glc mediated TOR-kinase antagonizes JA response

To further understand how the antagonistic interaction of Glc and MeJA occurs at the molecular level, we investigated the involvement of different components of Glc signal transduction in this response. Since TOR is a master regulator of Glc and energy signalling cascades (Sheen 2014) and that it regulates photosynthesis and phytohormone signalling pathways including JA signalling pathway (Dong *et al*. 2015), we wanted to assess the role of TOR-kinase in this response and for this we used *tor35-7 RNAi* line (Deprost *et al*. 2007). The RNAi line showed decrease in branching angle of LRs in response to independent and combined treatments of Glc and MeJA, as compared to WT seedlings (Fig. 2H-I; Fig. S5A). The *REGULATOR OF G-PROTEIN SIGNALLING* (*RGS1*)-dependent Glc pathway involves a G-protein signalling component, wherein RGS1 possibly acts as a Glc sensor (Urano *et al*. 2012b). To elucidate the role of *RGS1*-dependent Glc signalling components, Glc/JA regulation of branching angles in the *RGS1*-dependent signalling mutants *rgs1-1, rgs1-2, gpa1-3*, and *gpa1-4*. The RGS1-dependent signalling mutants showed no major change in LR angle as compared with their respective WT (Fig. S5B-C), suggesting that Glc does not utilise this pathway to regulate LR angle. In order to further substantiate the role of TOR signalling in this response, we treated Jas9-VENUS with Glc, alone and in combinations with MeJA and AZD-8055 (a potent inhibitor of TOR kinase activity) (Montané & Menand 2013). Intriguingly, we observed that Glc mediated accumulation of Jas9-VENUS was perturbed in seedlings treated with AZD (Fig. 2J-K; Fig. S6). Altogether, a TOR-dependent energy signalling mechanism is involved in the regulation of branching angle, and any perturbation to the mechanism leads to an altered JA response.

To further dissect the relationship between JA and Glc signalling at the whole genome transcriptome level, microarray analysis was employed. Most of the core JA signalling genes were down-regulated in the presence of 3%Glc and 3%Glc+MeJA when compared with 0%Glc+MeJA (Fig. S7). The microarray analysis thus, falls in agreement with our physiological and molecular data, confirming that antagonism operates between JA and Glc signalling pathways in regulating LR angle.

### Auxin transport and signalling lies downstream to MeJA-mediated modulation of branching angle

Recent reports have shed light on the role of PIN auxin efflux activity and the TIR1/AFB-Aux/IAA-ARF-dependent auxin signalling module for the establishment of GSA in young LRs (Rosquete *et al*. 2013, 2018; Roychoudhry *et al*. 2013, 2019). We, therefore, assessed the involvement of auxin transport and signalling to govern this physiological response. For this, the polar auxin transport inhibitor 1-N-naphthylphthalamic acid (NPA) was applied to the WT seedlings. NPA increased the Glc-induced branching angle (Fig. 3A). When applied in combination, NPA inhibited the MeJA response (Fig. 3A). To further investigate whether a functional auxin transport machinery is needed for MeJA to reduce branching angles, auxin transport defective mutants were examined for changes in branching angle. The auxin influx-defective mutant *auxin resistant 1 (aux1-7)* showed resistance to MeJA treatment and displayed expansive branching angles as compared to WT, with a greater percentage of LRs exhibiting angles >70° (Fig. 3B-C; Fig. S8A). The *lax3 (like <u>a</u>u<u>x</u>1 3)* mutant showed WT-like responses to MeJA treatment (Fig. S8B). Auxin-efflux defective mutants such as *ethylene insensitive root 1* (*eir1-1*), a mutant of *PIN2* and *multiple drug resistance 1* (*mdr1-1*), a mutant of *PGP1*, exhibited LRs with significantly broader angles compared with WT in the presence of Glc and in combination with MeJA (Fig. 3C-E; Fig. S8C). Since PIN3 is a major player in SII LR angle regulation (Rosquete *et al*. 2013), we examined the phenotype of *pin3-4* and observed no difference in LR angles when compared with WT (Fig. S8D). Other weak alleles of the *pinoid* mutants (*pin4-3* and *pin7-2*) exerted a WT like response to MeJA (Žádníkova *et al*. 2010) (Fig. S8D). The branching angle distribution of the auxin receptor mutant *transport inhibitor response 1* (*tir1-1)* had more LRs in the>70° category compared to WT (Fig. 4A-B; Fig. S8E). We also used *auxin resistant 1* (*axr1-3)* and *axr2-1*, a mutant of *IAA7*, which show constitutive downregulation of auxin responses (Leyser *et al*. 1993; Nagpal *et al*. 2000). Seedlings carrying the weak allele of *axr1-3* had near normal WT like responses to MeJA treatment (Fig. 4A, C; Fig. S8F). However, this response was completely abolished in *axr2-1* (Fig. 4A, C; Fig. S8F). Altogether, these results indicate that an intact auxin transport and signalling machinery is required for MeJA to set the angular growth of LRs.

**Figure 3.**
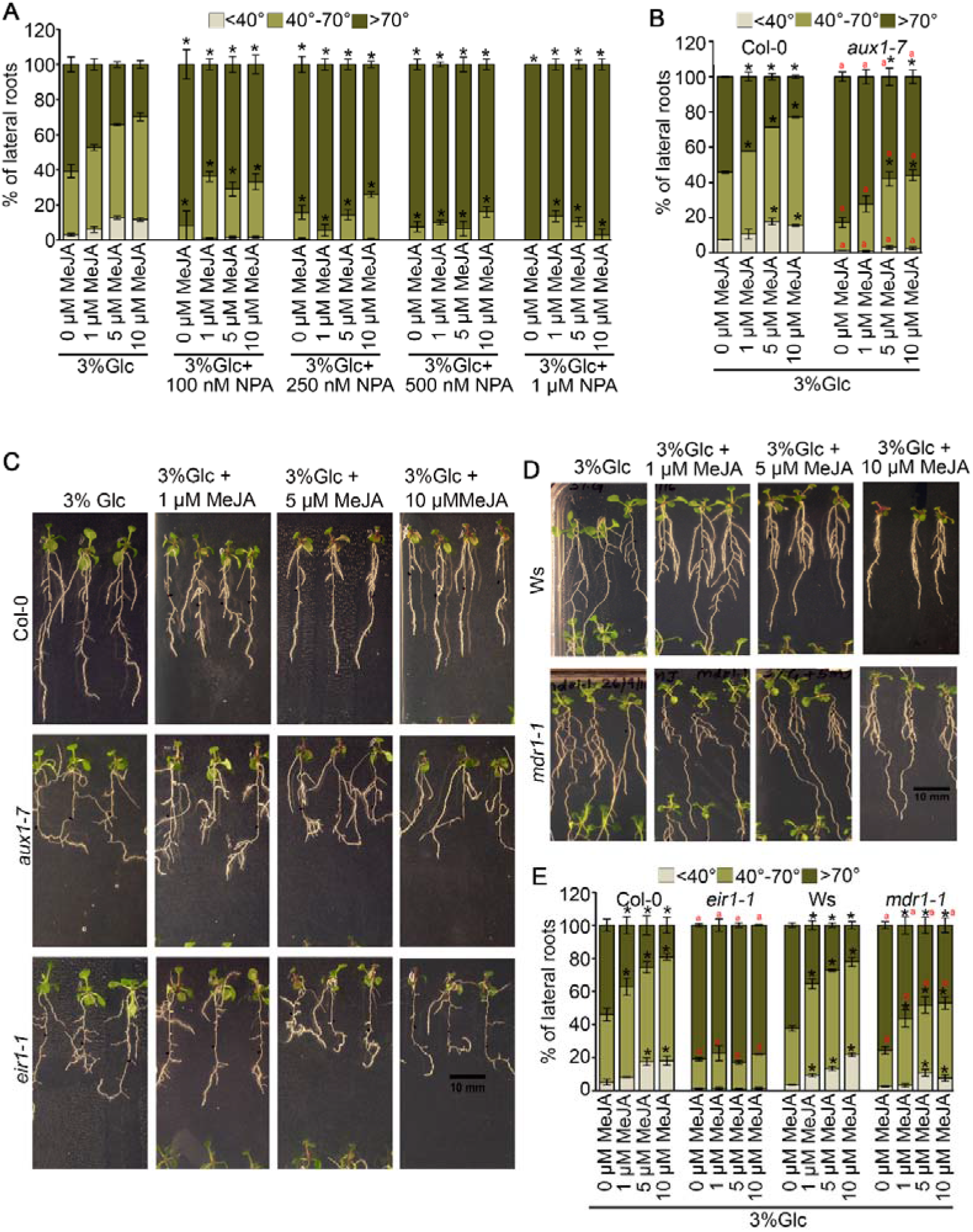
Components of auxin biosynthesis and transport lie downstream to MeJA-mediated branching angle. **(A)** Distribution of branching angle in Col-0 seedlings grown in 3%Glc alone or in combination with MeJA (10 μM) and different doses of NPA (100 nM to 1 μM). The graph represents the average of 4 biological replicates consisting of 25 seedlings and error bar represent SE. **(B)** Distribution of branching angle in seedlings of auxin influx defective mutant *aux1-7* grown in 3%Glc alone or in combination with different doses of MeJA. The graph represents the average of 3 biological replicates consisting of 25 seedlings and error bars represent SE. **(C)** Phenotype of light grown 12-day-old Arabidopsis Col-0, *aux1-7* and *eir1-1* seedlings grown in 3%Glc alone or in combination with different doses of MeJA. **(D)** Phenotype of light grown 12-day-old Arabidopsis Ws and *mdr1-1* seedlings grown in 3%Glc alone or in combination with different doses of MeJA. **(E)** Distribution of branching angle in seedlings of WT (Col-0 and Ws) and auxin efflux defective mutants *eir1-1* and *mdr1-1* grown in 3%Glc alone or in combination with different doses of MeJA. The graph represents the average of 5 biological replicates consisting of 25 seedlings and error bars represent SE. 5-day-old 1/2 MS grown Col-0 and mutant seedlings were transferred to treatment media and phenotypes were analysed at 12^th^ day. Asterisks indicate a significant difference in the studied parameter (P <0.05, paired two-tailed student’s t-test; * control vs treatment and ‘a’ WT vs mutant).

**Figure 4.**
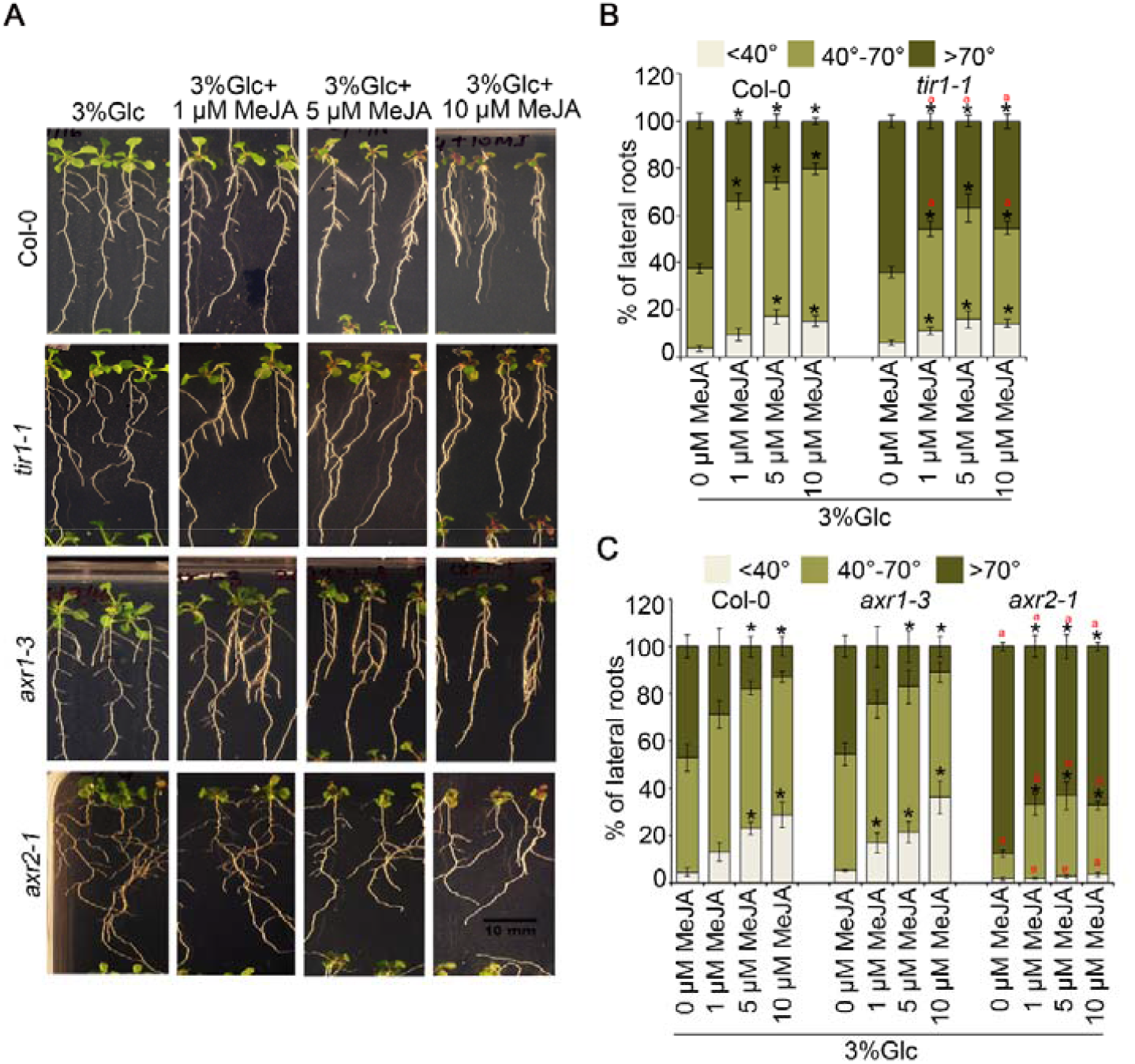
Components of auxin signalling lie downstream to MeJA-mediated branching angle. **(A)** Phenotype of light grown 12-day-old Arabidopsis Col-0, *tir1-1, axr1-3* and *axr2-1* seedlings grown in 3%Glc alone or in combination with different doses of MeJA. **(B-C)** Distribution of branching angle in seedlings of auxin signalling defective mutants *tir1-1, axr1-3* and *axr2-1* grown in 3%Glc alone or in combination with different doses of MeJA. The graph represents the average of 4 biological replicates consisting of 25 seedlings and error bars represent SE. 5-day-old 0.5X MS grown Col-0 and mutant seedlings were transferred to treatment media and phenotypes were analysed at 12^th^ day. Asterisks indicate a significant difference in the studied parameter (P <0.05, paired two-tailed student’s t-test; * control vs treatment and ‘a’ WT vs mutant).

### MYC2 controls the transcription of *CYP79B2, LAZY2* and *LAZY4*

A former report has shown that jasmonate mediates the regulation of auxin biosynthesis (Sun *et al*. 2009). *CYP79B2*, a cytochrome P450 mono-oxygenase that forms indole-3-acetaldoxime and acts as a precursor for auxin biosynthesis (Zhao *et al*. 2002). Additionally, other molecular components are involved in modifying plant architecture, including the *LAZY* gene family, which controls both root and shoot gravitropism (Taniguchi *et al*. 2017; Yoshihara & Spalding 2017). *LAZY2 (DRO3/NGR1), LAZY3 (DRO2/NGR3)*, and *LAZY4 (DRO1/NGR2)* have been reported to cause auxin redistribution and transport in *Arabidopsis* LRs; thus, leading to an overall vertical root architecture (Taniguchi *et al*. 2017; Yoshihara & Spalding 2017). Mutants defective in *LAZY2* and *LAZY4* have agravitropic LRs (Taniguchi *et al*. 2017; Yoshihara & Spalding 2017), prompted us to study jasmonate effects on the LAZY2 and LAZY4-mediated LR angles. Given the established role of JA in transcriptionally activating *CYP79B2*, and that MYC2 is the master regulator of many JA responses, it is reasonable to hypothesize that MYC2 might control the transcription of these genes, enhancing LR response to gravity. To test this hypothesis, we examined the MYC2-depenedent expression of *CYP79B2, LAZY2* and *LAZY4*. As shown in Figure 5A-C, *CYP79B2, LAZY2* and *LAZY4* transcript levels were downregulated in *myc2myc3myc4*. To understand how MYC2 regulates the expression of these genes, we scanned the promoters of *CYP79B2, LAZY2* and *LAZY4*, and found G- and E-box elements. Using chromatin immunoprecipitation-quantitative PCR (ChIP-qPCR), we found that *CYP79B2, LAZY2*, and *LAZY4* promoters were highly enriched with MYC2 protein in *35S::MYC2-GFP* when compared to WT (Fig. 5D-F). We also examined the binding of MYC2 to the promoters of *ORA59*, which is a known MYC2 target and served as a positive control for our experiments (Zhai *et al*. 2013). *ATXR6* was used as a negative control because it does not possess any MYC2 binding sites in its promoter region (Fig. 5D-F). The results suggest that MYC2 binds and controls the transcription of genes that can influence the vertical orientation of LRs.

**Figure 5.**
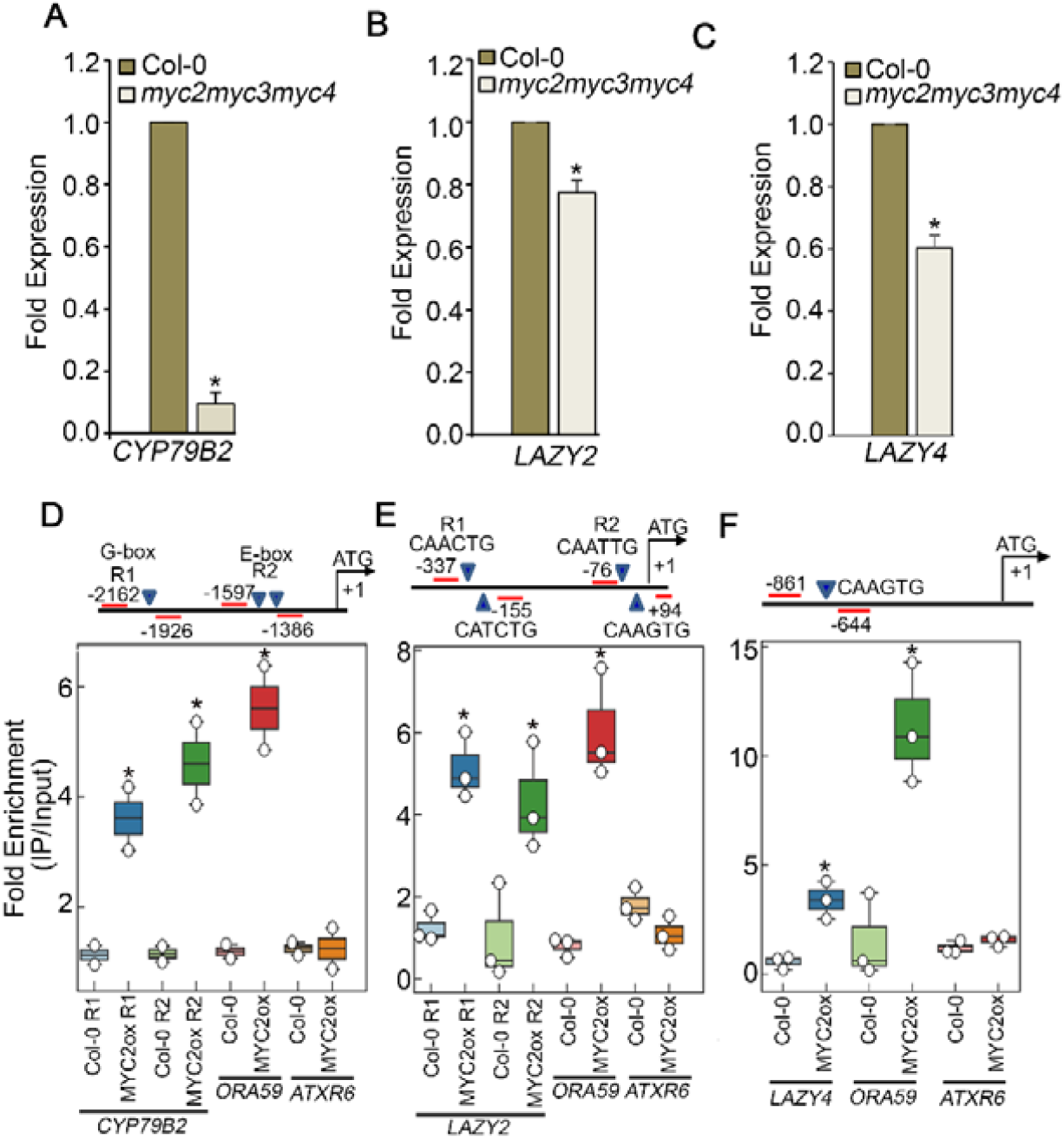
Transcriptional regulation of *CYP79B2, LAZY2* and *LAZY4* by MYC2. **(A-C)** RT-qPCR showing the expression of *CYP79B2, LAZY2* and *LAZY4* in 1/2 MS media in Col-0 and *myc2myc3myc4* seedlings. The graph represents the average of 3 biological replicates consisting of 25 seedlings and error bars represent SE. **(D)** ChIP-qPCR showing the enrichment of *CYP79B2* promoter fragments with 35S::MYC2::GFP in all two regions. The plot shows average of 2 biological replicates. **(E)** ChIP-qPCR showing the enrichment of *LAZY2* promoter fragments with MYC2 in all two regions. **(F)** ChIP-qPCR showing the enrichment of *LAZY4* promoter fragments with MYC2. Fold enrichment of promoter fragments was calculated by comparing samples treated without or with anti-GFP antibody. Untransformed Col-0 was taken as a negative genetic control. The plot shows average of 3 biological replicates. *ORA59* and *ATXR6* were used as positive and negative controls, respectively. Asterisks indicate a significant difference in the studied parameter. Error bars represent SE. (P <0.05, paired two-tailed student’s t-test; * WT vs overexpression).

### MeJA controls the direction of auxin transport that may regulate LR angle

There are various factors that control the GSA of LRs via the regulation of auxin flow from LR tips (Claudia-Anahi’ Pèrez-Torres *et al*. 2008; Roychoudhry *et al*. 2017; Taniguchi *et al*. 2017). These reports led us to investigate whether JA signalling influences auxin distribution at the Stage II (SII) LR tips. For this, we analysed the effect of MeJA on *DR5::GFP* expression. Most of the 1/2 MS-treated LRs displayed near symmetrical *DR5::GFP* expression with a faint green signal streak towards the lower side of the LR tip (Fig. 6A). However, after the addition of MeJA, the *DR5::GFP* expression started to disappear from the upper side and the green signal became more apparent on the margins of the lower side (Fig. 6A). These observations suggested that JA signalling directs the flow of auxin towards the lower side, which might lead to vertical orientation of SII LRs. Under control conditions (1/2 MS), the expression of *PIN3::PIN3-eGFP* was strongly expressed in the columella cells of SII LRs. However, in seedlings treated with MeJA, there was an expansion of GFP signal towards the stelar region of SII LRs but the signal intensity was very low. Also, there weren’t any changes in the differential distribution of PIN3::PIN3-GFP upon MeJA treatment. Upon quantification of the signal intensity, we did not find any significant difference in its expression between control and MeJA treatment, suggesting that JA signalling might not involve PIN3 mediated auxin redistribution (Fig. S9A-B). In addition to PIN3 activity in the columella, the action of PIN2 in the epidermis is required to drive the basipetal flow of auxin away from the root tip (Roychoudhry *et al*. 2019). We, therefore, studied *PIN2::PIN2-eGFP* expression in LRs growing at their GSA to check for any differential expression of PIN2 that might contribute to the regulation of growth angle. The *PIN2::PIN2-eGFP* signal intensity was strong and intact in 1/2 MS-treated seedlings. Also, there was not any difference in PIN2 expression between upper and lower halves of LRs (Fig. 6B). However, we observed that MeJA diminished PIN2 redistribution in the upper epidermal cells of the LR (Fig. 6B). Additionally, we found that the length of the 1^st^ two upper epidermal cells of LR treated with MeJA was longer than that in the control (Fig. 6B-C). It has been previously revealed that differential elongation is a major factor controlling LR gravitropism (Rosquete *et al*. 2013) and therefore, these results suggest that the JA signalling via PIN2 plays a potential role in differential cell elongation, leading to vertical orientation of LRs.

**Figure 6.**
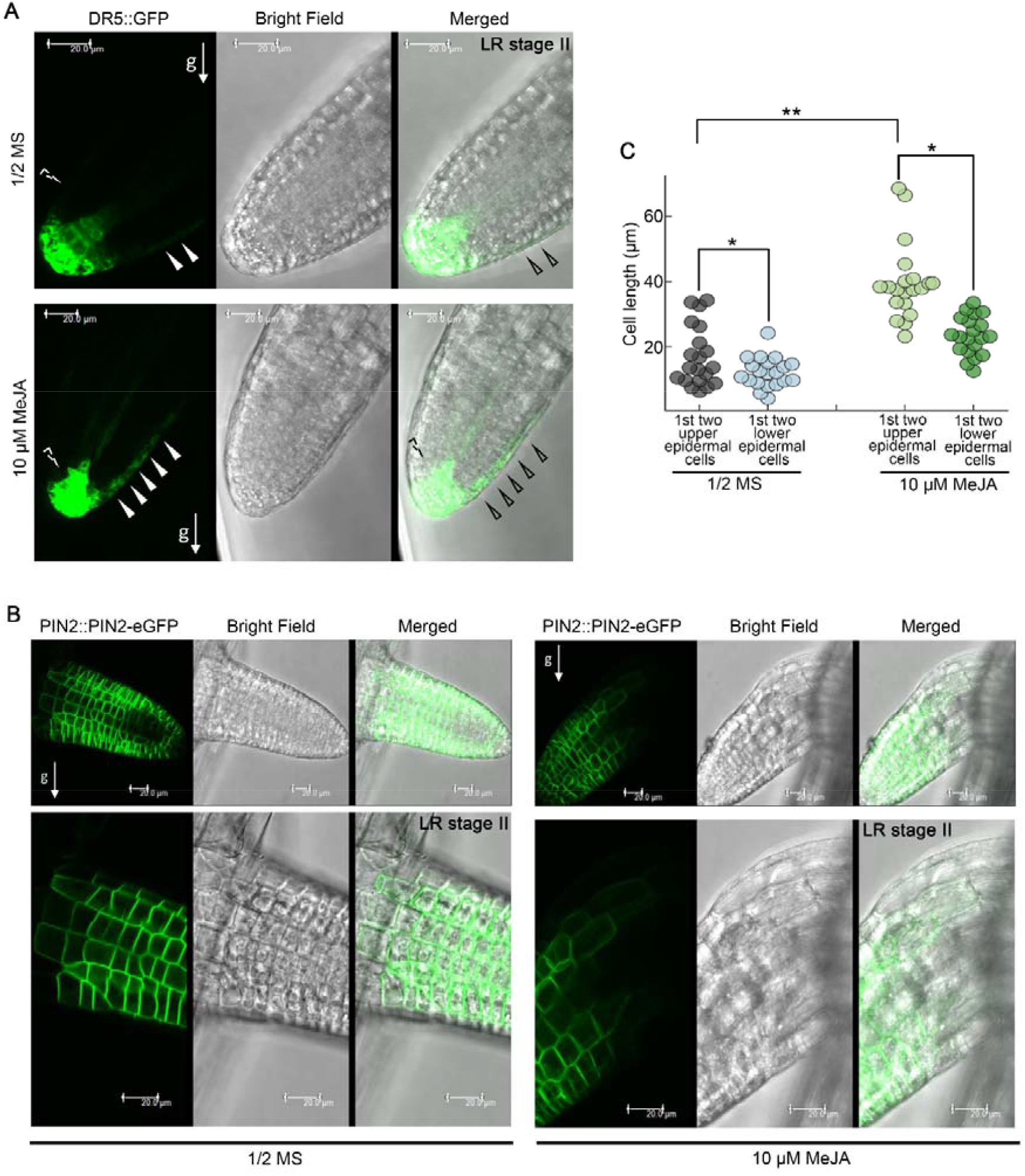
MeJA controls lateral auxin redistribution in LRs. **(A)** Stage II LR of DR5::GFP seedling treated with 1/2 MS and in combination with MeJA (10 μM). 5-day-old 1/2 MS grown Arabidopsis DR5::GFP seedlings were transferred to 1/2 MS alone or in combination with MeJA. At 12^th^ day, stage II lateral root were assessed under confocal microscope. Scale bar: 20 μm. The data was repeated three times, with 8-9 roots imaged every time. Solid white arrowheads depict strong GFP signal at the LR tip and dotted white arrowheads depict reduced GFP signal. **(B)** Upper panel showing stage II LR of PIN2::PIN2-eGFP seedlings treated with 1/2 MS and in combination with MeJA (10 μM). 5-day-old 1/2 MS grown Col-0 seedlings were transferred to treatment media. Lower panel showing enlarged view of 1/2 MS and MeJA treated PIN2::PIN2-eGFP LR with diminished PIN2 activity from the upper epidermal cell profile. **(C)** The graph represents the cell length of 1^st^ two upper and lower epidermal cells of LRs treated with 1/2 MS alone and in combination with 10 μM MeJA. The data shown is an average of two biological replicates. (P <0.05, paired two-tailed student’s t-test; * upper and lower epidermal cell profile; ** control vs treatment). The data was repeated two times with 10-12 roots imaged.

### MeJA integrates environmental cues into angular growth of LRs

Our data highlights a role for JA signalling in modulating angular growth of LRs. Also, light can modify the GSA of organs (Digby & Firn 2002; Roychoudhry *et al*. 2017). To investigate whether light affects MeJA mediated branching angle, 5-day-old light-grown WT and *myc2myc3myc4* seedlings were transferred to 0.5%Glc, 0.5%Glc+MeJA, 3%Glc, and 3%Glc+MeJA under long-day regime (16 hours light/8 hours dark) and under continuous dark conditions. In the dark, seedlings showed elongated hypocotyls and significantly more horizontally placed LRs than those grown in light conditions (Fig. S10A-B). In both WT and *myc2myc3myc4*, the effect of 3%Glc was more prominent in dark conditions, with LRs displaying >70° angles. MeJA treatment did not decrease the angles in dark conditions. Also, LRs of *myc2myc3myc4* showed wider angles with MeJA treatment in dark conditions as compared to light (Fig. S10A-B), thus suggesting that light is required for an optimal MeJA response. Next, to investigate whether light affects auxin distribution in controlling branching angle, we treated WT and *aux1-7* seedlings in light and dark regimes and observed that WT seedlings showed broader angles in dark as compared to light (Fig. S11). Interestingly, we found that *aux1-7* showed broader branching angle under both light and dark conditions, suggesting that light modulates auxin distribution and ultimately affects branching angle. We also observed that *aux1-7* was less responsive to MeJA treatment under both light and dark conditions (Fig. S11). Collectively, these results suggest that light and JA signalling are prerequisite to influence auxin distribution in governing branching angle.

Light perception and signalling intersect with the action of jasmonates to affect plant development and defence (Dombrecht *et al*. 2007). We, therefore, explored the involvement of phytochrome (PHY) mediated light signalling in regulating this response. Photoreceptor single mutants *phyA-201* and *phyB-5* showed broader angles when compared with WT, whereas the *phyA201phyB5* showed a much stronger response and exhibited completely altered Glc and MeJA sensitivity, with LRs showing horizontal angles at all concentrations (Fig. 7A-B). Hence, *PHYA* and *PHYB* act redundantly in controlling the phenotype. To further substantiate the role of light signalling in regulating this response, we investigated the involvement of *HY5*, a major downstream positive regulator of phytochrome signalling (Gangappa & Botto 2016). The *hy5-1* mutant seedlings, which are already known to show wider LR angles, (Oyama, Shimura & Okada 1997) displayed horizontally positioned LRs upon JA treatment as compared to the WT (Fig. 7A-B). Previous reports of ChIP-seq data showed that the binding of HY5 to the promoter of *LIPOXYGENASE3 (LOX3*), a JA biosynthetic gene (Lee *et al*. 2007). To further confirm whether light signalling is involved in maintaining an optimal JA response by increasing JA biosynthesis, we treated WT and *hy5-1* seedlings in long day and continuous dark conditions for 6 days. We also shifted 6-day-old continuous dark grown WT and *hy5-1* seedlings to light for 6 hours and checked for the expression of *LOX3*. Indeed, we found a significant increase in the expression of *LOX3* in long-day treatment when compared to total darkness (Fig. 7C). Also, WT seedlings exposed to a 6 hours dark to light transition exhibited enhanced LOX3 expression (Fig. 7D). In contrast, *LOX3* expression was significantly downregulated in *hy5-1* seedlings, suggesting that the response is *HY5*-dependent. Collectively, the above data suggest that JA signalling integrates environmental factor, such as light, into GSA establishment of LRs.

**Figure 7.**
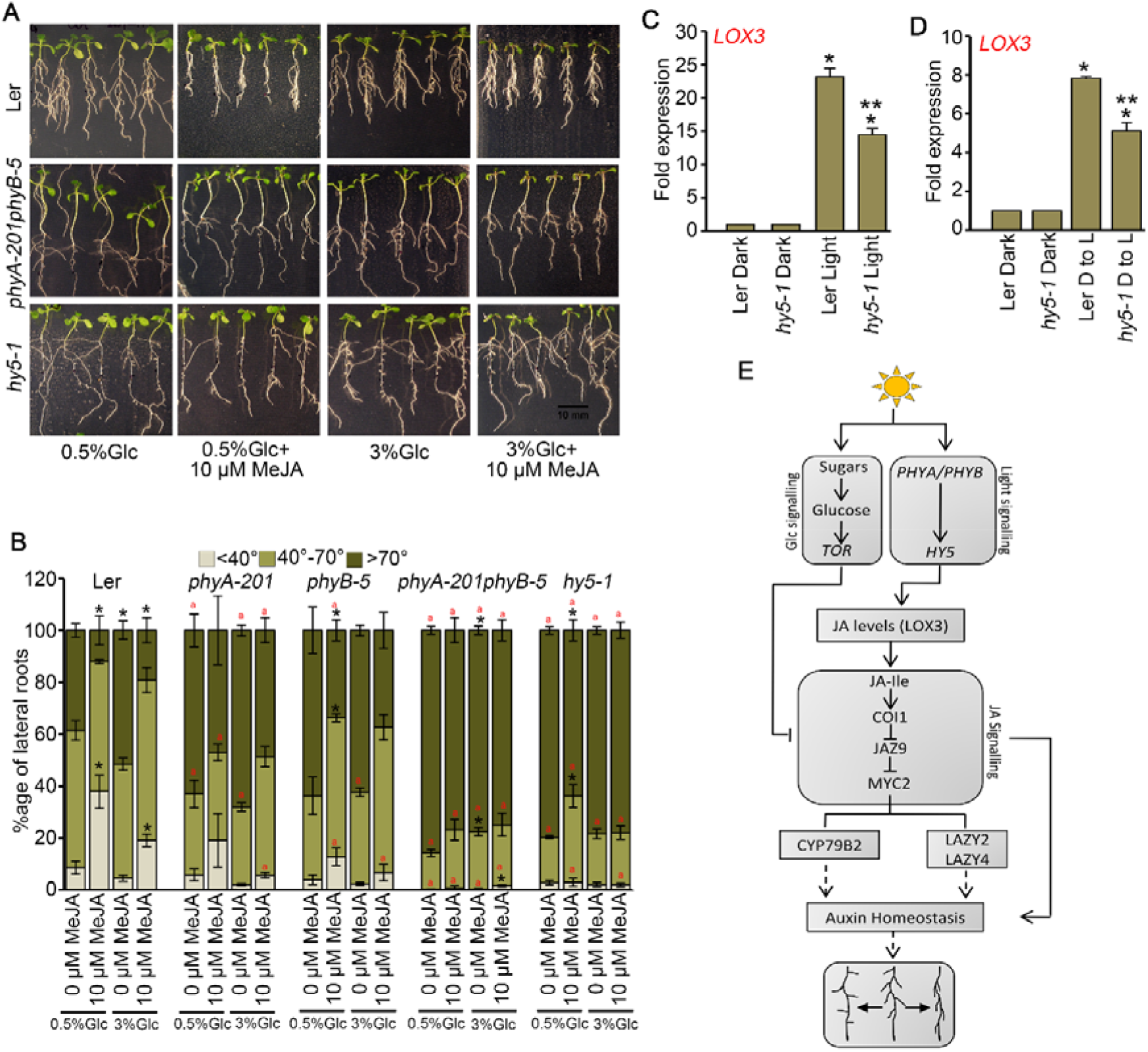
Light modulation of Arabidopsis root branching angle. **(A)** Phenotype of light grown 12-day-old Arabidopsis Col-0, *phyA201phyB5* and *hy5-1* seedlings treated with 0.5%Glc and 3%Glc alone or in combination with 10 μM MeJA. **(B)** Distribution of branching angle in seedlings of Col-0 and light signalling mutants *phyA-201, phyB-5, phyA201B5* and *hy5-1* grown in different concentrations of Glc (0.5%, 3%) alone or in combination with MeJA (10 μM). 5-day-old 1/2 MS grown Ler and mutant seedlings were transferred to treatment media and phenotypes were analysed at 12^th^ day. The graph represents the average of 3 biological replicates consisting of 25 seedlings and error bars represent SE. Asterisks indicate a significant difference in the studied parameter (P <0.05, paired two-tailed student’s t-test; * control vs treatment and ‘a’ WT vs mutant). **(C)** RT-qPCR showing the expression *LOX3* in 1/2 MS media under continuous light and dark for 6 days in Ler and *hy5-1* seedlings. **(D)** RT-qPCR showing the expression *LOX3* in 1/2 MS media under dark (D) and dark (D) to light (L) transition for 6 hours in Ler and *hy5-1* seedlings. The graph represents the average of 3 biological replicates, error bars represent SE. Asterisks indicate a significant difference in the studied parameter (P <0.05, paired two-tailed student’s t-test; * control vs treatment and ** WT vs mutant). (E) Model depicting interaction between light, Glc, JA and auxin signalling in regulating branching angle of Arabidopsis LRs.

## Discussion

The angular growth of LRs (GSA) represents an important element in the adaptability of the root system to its environment. Both nutrient and hormonal signals act locally to regulate GSA (Bai *et al*. 2013; Rosquete *et al*. 2013, 2018; Roychoudhry *et al*. 2013, 2017, 2019; Trachsel *et al*. 2013). The net effect of this adaptive response is an increase in the surface area of the plant root system for resource capture (e.g. horizontal LRs for phosphorus uptake), or the securing of anchorage (Lynch & Brown 2001; Trachsel *et al*. 2013). Similarly, shoots with vertical lateral branches show enhanced efficiency of light capture, allowing higher density planting and higher yields (Sakamoto *et al*. 2006; Vriet, Russinova & Reuzeau 2012). The main aim of this study is to advance our understanding of GSA regulation and to elucidate interconnections between environmental and hormonal signals in fine-tuning the branching angle of *Arabidopsis* roots.

In this work, we demonstrated that MeJA reduced the branching angle of roots in a dose-dependent manner, thereby resulting in an overall vertical orientation. Analysis of mutants defective in JA biosynthesis and signalling revealed that an intact JA biosynthesis and signalling is prerequisite to bring about changes in the branching angle. The branching angles of *jar1-11, jin1-9*, and *myc2myc3myc4* mutants were defective in perceiving endogenous JA, even under control conditions (1/2 MS), suggesting that, in nature, JA signalling plays an important developmental role in governing root branching angle. The weaker phenotype in *jin1-9* compared to that of *myc2myc3myc4* suggested that *MYC3* and *MYC4* act additively with *MYC2* in regulating this response. Since *MYC3* and *MYC4* are weakly expressed in the roots of young seedlings unlike *MYC2* (Fernández-Calvo *et al*. 2011), and *MYC2* is the major regulator of many JA governed biological responses (Lorenzo, Chico, Sánchez-Serrano & Solano 2004; Dombrecht *et al*. 2007), we postulate that this response is majorly mediated by *MYC2*.

Apart from affecting various parameters of RSA (Gupta, Singh, Mishra, Kushwah & Laxmi 2009; Gupta *et al*. 2015; Mishra *et al*. 2009; Singh *et al*. 2014a b), sugars can also influence the gravitropic behaviour of lateral organs (Willemoes *et al*. 1988). Previous studies have shown that 3%Glc causes more production of LRs and root growth and is therefore, considered optimal for overall plant growth and development (Mishra *et al*. 2009; Singh *et al*. 2014a; Gupta *et al*. 2015). In this study, 3%Glc caused a significant shift in the branching angle towards a more horizontal orientation. This effect of Glc on growth can favour plant propagation because it allows plants to explore adjacent territories. The diagravitropic growth of LRs upon Glc treatment was consistent with a report by Willemoës *et al*. (1998), suggesting that sugars are responsible for maintaining a non-vertical angular growth of organs (Willemoes *et al*. 1988). There are also reports citing crosstalk between sugars and GA_3_ to regulate growth direction of organs (Montaldi 1969; Willemoes *et al*. 1988). Compared with well-studied interactions between Glc and other hormones, relatively little is known about Glc and JA (Song *et al*. 2017; Vleesschauwer *et al*. 2017; Guo *et al*. 2018). Most of these studies demonstrate an antagonistic interaction between these two signalling pathways in both dicots and monocots (Song *et al*. 2017; Vleesschauwer *et al*. 2017). Moreover, transcriptomic analysis revealed the regulation of JA signalling pathway by TOR-kinase (Dong *et al*. 2015). In our study, microarray analysis, immunoblot assays and physiological studies have confirmed that Glc and JA are two crucial signals that work antagonistically via TOR kinase pathway to regulate the branching angle of *Arabidopsis* roots. This observation falls in agreement with the previous report citing antagonistic interaction between JA and TOR kinase (Song *et al*. 2017). Transcriptome analysis of rice cells treated with the TOR-specific inhibitor rapamycin revealed that TOR, apart from dictating transcriptional reprogramming of extensive gene sets involved in central and secondary metabolism, cell cycle, and transcription, also suppresses many defence-related genes and antagonizes the action of JA (Vleesschauwer *et al*. 2017). We hypothesize that this antagonism occurs in nature in order to fine-tune the response and achieve the optimum angle required for growth. However, more detailed molecular dissection is necessary to obtain a deeper understanding of this crosstalk and its regulators.

Of all the plant hormones, auxin plays a central role in LR GSA control. Transient expression of *PIN3* with strong repression of *PIN4* and *PIN7* in young LRs limits auxin redistribution and hence explains the reduced gravitropic competence of laterals (Rosquete *et al*. 2013, 2018). JAs contribute to the regulation of transport of IAA by inducing the expression of *PIN1* and *PIN2* (Sun *et al*. 2009) and modulate the accumulation of PIN2 in the plasma membrane and its recycling via endocytosis, in a dose-dependent manner (Sun *et al*. 2011). In light of the above reports claiming recent interconnections between JA and auxin signalling, and the modulation of JA homeostasis and signal transduction mimicking auxin effects on root development, we show that MeJA requires auxin machinery to control branching angle. Our results suggest that genetic disruption of auxin transport and signalling nullifies the effect of MeJA-mediated control of branching angle. Additionally, due to differences in the phenotypic strength of the mutants, MDR1-mediated auxin transport and TIR1-AXR1-mediated auxin signalling show much fewer effects than AUX1-PIN2 and AXR2-mediated transport and signalling (Tiryaki & Staswick 2002; Quint, Barkawi, Fan, Cohen & Gray 2009). JA signalling via MYC2 also induces the transcription of *CYP79B2, LAZY2* and *LAZY4*. We suppose that this transcriptional activation might promote auxin biosynthesis and its transport and redistribution, ultimately leading to vertical angles. Also, the modulation of the asymmetric auxin transport in LR columella cells as observed by DR5::GFP expression, diminished PIN2 activity from the upper epidermal cell profile, and the differential cell elongation caused by MeJA may correlate with the vertical orientation of the LRs.

Light is one of the many diverse signals that changes the sugar/energy status in plants. Studies till date have given tremendous insights into how light not only drives photosynthesis but also acts as an environmental cue that informs plants about their environment. Previous literature have stated that light can increase root gravitropism by influencing the direction of polar auxin transport (Buer & Muday 2004; Laxmi, Pan, Morsy & Chen 2008; Sassi *et al*. 2012). Consistently, our results demonstrate that light acts as a stimulus to change auxin distribution to modulate branching angle of *Arabidopsis* LRs. Also, the exogenous application of MeJA could not restore the vertical orientation of branching angle in *aux1-7* neither in dark nor in light, further suggesting that auxin distribution is indispensable for MeJA modulation of branching angle in light. Light signalling also influences auxin transport in the roots by vacuolar degradation of PIN proteins (Korbei & Luschnig 2013) and is essential for regulation of root gravitropism. Light via COP1/CSN complex controls the vacuolar targeting of PIN2-GFP (Laxmi *et al*. 2008; Sassi *et al*. 2012).

Previous reports have demonstrated that light quality governs the gravitropic behaviour of lateral organs (Kiss *et al*. 2002). *Arabidopsis* LRs showed negative phototropism in the presence of white light, and positive phototropism in response to red light (Kiss *et al*. 2002). In the present study, we observed that light, acting via the redundant functions of PHYA-PHYB and HY5 induced a more vertical LR orientation, supporting results reported by Kiss *et al*. (2002). Light quality also influences formation of lateral root primordia by accumulation of HY5. Also, HY5-mediated light signalling influences root development by regulating auxin transport and signalling (van Gelderen *et al*. 2018). In our experiments, exogenous application of MeJA could not induce vertical branching angles of LRs in dark conditions. This suggests that light is prerequisite to MeJA-induced changes in branching angle. We assume that the alteration in the angle is not only achieved by the biosynthesis of JA, but also by the influence of another factor or signalling event that is currently unknown. ChIP-seq data showed the binding of *HY5* at the promoter of *LOX3* (Lee *et al*. 2007). Consistent with this observation, we found decreased expression levels of *LOX3* in *hy5-1* under light conditions as well as dark to light transition. This suggests a likelihood that HY5 regulates JA levels in light. Other studies have shown that light environment and circadian rhythms are crucially involved in modulating plant responses to JA biosynthesis (Radhika, Kost, Mithofer & Boland 2010; Goodspeed, Chehab, Min-Venditti, Braam & Covington 2012). JA levels have been shown to rise during the day, reaching a maximum at midday, and then declining again in the afternoon (Goodspeed *et al*. 2012). MYC2 protein levels were also shown to rise during the day and under continuous light conditions (Shin, Heidrich, Sanchez-Villarreal, Parker & Davis 2012). Moreover, ChIP-seq analyses show the binding of HY5 on the promoter of *MYC2* (Lee *et al*. 2007). Considering previous literature and our current findings, we show that light either directly or indirectly can dramatically affect plant root growth and development. We have, therefore, in this study have unraveled the interconnection of light signalling and its modulation by glucose (a downstream product of light driven photosynthesis) and phytohormones like JA and auxin in controlling root branching angle.

Based on our investigation and previous findings, we propose a testable model in Figure 7E. We hypothesize that light works via two branches to optimize the branching angle of *Arabidopsis* LRs. Contrary to the general notion that light and sugars should have the same influence on developmental outputs, there are many reports that suggest that light and Glc signalling have opposite effects on the growth and development of *Arabidopsis* seedlings (Moore 2003; Eckstein, Zieba & Gabryś 2012). Light working via phytochrome signalling (*PHYA/PHYB-HY5*) increases JA levels, which ultimately lead to the activation of JA signalling and this physiological response is shown to be mainly mediated by *MYC2*. Our results also suggest that auxin works further downstream as auxin transport and signalling mutants displayed abrogated responses to JA regulation of branching angle. Additionally, MYC2 regulates the expression of *CYP79B2, LAZY2* and *LAZY4* by binding to their promoters that might promote auxin biosynthesis and its transport and redistribution, ultimately leading to vertical angles. Glc, on the other hand, is produced by photosynthesis promotes radial expansion of the root architecture, antagonistically interacting with JA signalling via the *TOR*-mediated pathway and affecting the accumulation of JAZ9 to regulate this developmental process.

We hypothesize that under normal conditions, roots grow in the soil without light. Upon light exposure, the stress hormone JA is induced that further regulates auxin transport and signalling to accelerate the vertical growth of LRs to avoid light. However, under sufficient energy (3%Glc), roots do not feel stressed and hence, do not seem to avoid light and grow horizontally. Though, horizontal angles favour plant propagation and explore adjacent territories better but do not play a significant role in root anchorage (due to less anchorage force) (Ennos 2000; Mickovski & Ennos 2003), Similarly, steeper LRs show less soil column weight and hence low uprooting resistance. Thus, to maintain a balance, MeJA and Glc signalling act in concert to maintain an optimal angle where LRs act as guy ropes, holding the vertical root in position that not only favours plant propagation but also provides maximum anchorage to prevent uprooting of plants (Ennos 2000). Apart from the root architecture, signals from the environment and endogenous cues act together to optimize growth angles of lateral branches. Depending upon the strength of the signal, the angle of the laterals is decided. The present study in *Arabidopsis* can be used for crop improvement strategies that can lead to crop species with improved water and nutrient acquisition capability and hence is of broad relevance to plant and crop scientists. Future work will be directed at identifying substantial mechanistic insights into hormonal-environmental crosstalk and characterizing novel molecular components that are involved in shaping the overall root architecture.

## Acknowledgements

The authors would like to thank Dr. Aditi Gupta for her advice and discussions and Ms Harshita Bharti Saksena for reading the manuscript. The authors acknowledge NIPGR Confocal Facility for their assistance, Prof. Philippe Reymond for providing seeds of *myc2myc3myc4*, Dr. Nam-Hai Chua for providing seeds of *35S-MYC2-GFP* and Dr. Laurent Laplaze for providing seeds of *p35S::Jas9-N7-VENUS*. The authors are thankful to DBT-eLibrary Consortium (DeLCON) for providing access to e-resources. This work was financially supported by the Core Grant from the National Institute of Plant Genome Research to A.L., University Grant Commission, Government of India and Department of Biotechnology, Government of India. M.S. acknowledges University Grant Commission, Government of India for research fellowship, M.S. acknowledges Department of Biotechnology, Government of India and MJK acknowledges Department of Science and Technology (INSPIRE Faculty Programme Grant IFA18-LSPA110).

## Author Contributions

M.S. and A.L. conceived and designed the experiments. M.S. performed physiology and microarray. M.S, M.S^#^ and M.J.K performed confocal experiments. M.S and M.S^#^ performed real time assays, western blot and ChIP-qPCR. M.S. wrote the article. M.S^#^ and M.J.K. assisted in microarray analysis and preparing the manuscript; A.L. supervised and complemented the article.

## Conflict of Interest statement

The authors declare no conflict of interest.

## Supplementary data

**Figure S1. MeJA regulation of branching angle of *Arabidopsis* roots. (A and B)** Average branching angle of 12-day-old Col-0 and JA signalling mutants grown in different doses of MeJA. The data represents the average of 4 biological replicates consisting of 25 seedlings and error bars represent SE. **(C)** Distribution of branching angle of Col-0 and different *jaz* seedlings grown in different doses of MeJA. The data represents the average of 3 biological replicates consisting of 25 seedlings and error bars represent SE. 5-day-old 1/2 MS grown Col-0 and mutant seedlings were transferred to treatment media and phenotypes were analysed at 12^th^ day. Asterisks indicate a significant difference in the studied parameter (P <0.05, paired two-tailed student’s t-test; * control vs treatment and WT vs mutant).

**Figure S2. MeJA regulation of branching angle of *Arabidopsis* roots. (A)** A representative image of 20 days after germination (DAG) of Col-0 and JA biosynthesis and signalling mutants *jar1-11* and *myc2myc3myc4* grown in 1/2 MS medium in cylindrical tubes. **(B)** Average branching angle of 20 DAG of Col-0, *jar1-11* and *myc2myc3myc4* grown in 1/2 MS medium in cylindrical tubes. The data represents images from 2 biological replicates, each having 5 plants (P <0.05, paired two-tailed student’s t-test; * WT vs mutant).

**Figure S3. MeJA-Glc regulation of branching angle of *Arabidopsis* roots. (A)** Average branching angle of 12-day-old Col-0 seedlings grown in different concentrations of Glc (0.5%, 3%) and in combination with 10 μM MeJA. The data represents the average of 7 biological replicates consisting of 25 seedlings and error bars represent SE (P<0.05, Student’s t-test; * control (0.5%Glc) vs treatment. (B) Phenotype of Col-0 seedlings grown in different concentrations of Glc (0.5%; 28 mM and 5%Glc; 278 mM) and in combination with 10 μM MeJA. (C) Distribution of branching angle of seedlings of Col-0 grown in different different concentrations of Glc (0.5%, 3% and 5%) and in combination with MeJA. The data represents the average of 4 biological replicates consisting of 25 seedlings and error bars represent SE. Asterisks indicate a significant difference in the studied parameter (*P*<0.05,Student’st-test;*control (0.5%) vs treatment).

**Figure S4. MeJA-Glc regulation of branching angle of *Arabidopsis* roots. (A)** 3 individual biological replicates of Jas9-VENUS seedlings treated with increasing concentrations of Glc (0%Glc, 0.5%Glc, 3%Glc) along with 10 μM MeJA. **(B)** RT-qPCR showing the expression JA signalling genes (*JAZ1, JAZ2, JAZ3, JAZ6, JAZ9, AOS, MYC2, ERF1* and *ORA59*) in Col-0 seedlings grown in 0%Glc and 0%Glc+ 10 μM MeJA. 5-day-old 1/2 MS grown Col-0 seedlings were first starved for 24 hrs and transferred to liquid treatment media containing 10 μM MeJA and analysed after 3hrs. The data represents the average of 4 biological replicates consisting of 25 seedlings and error bars represent SE (P <0.05, paired two-tailed student’s t-test; * control (0%Glc) vs treatment (0%Glc+MeJA).

**Figure S5. Role of different components of Glc signalling in regulation of branching angle of *Arabidopsis* roots. (A)** Average branching angle of 12-day-old Col-0 and Glc-dependent TOR signaling mutant *tor35-7RNAi* grown in different concentrations of Glc (0.5%, 3%) and in combination with MeJA. The data represents the average of 4 biological replicates consisting of 25 seedlings and error bars represent SE. (*P* <0.05, Student’s t-test; * control vs treatment ** 0.5%Glc vs 3%Glc+10MeJA and wild-type vs mutant). **(B and C)** Distribution of branching angle in seedlings of Col-0 and RGS-dependent signalling mutants *rgs1-1, rgs1-2, gpa1-3* and *gpa1-4* grown in different concentrations of Glc in combination with MeJA. The data represents the average of 4 biological replicates consisting of 25 seedlings and error bars represent SE.

**Figure S6. Glc regulates branching angle of LRs via TOR-mediated signalling pathway**. 3 individual biological replicates of Jas9-VENUS seedlings treated without and with Glc (3%Glc) along with 10 μM MeJA, 10 μM AZD and a combination of 10 μM MeJA and 10 μM AZD (related to Figure 2J, K).

**Figure S7. Heat map of JA signalling genes upon treatment with 0%Glc+10μM MeJA, 3%Glc and 3%Glc+10μM MeJA**. The fold change value of genes (+/-1.5 fold) treated with 0%Glc+10μM MeJA was considered as 1. The fold change values of genes in 3%Glc and 3%Glc+ 10 μM MeJA was normalised (as % change) with those of 0%Glc+10μM MeJA. * indicates expression values of genes that are significantly altered. The data is the average of 2 biological replicates. Heat map was generated using MeV.

**Figure S8. Role of auxin machinery in controlling JA-mediated branching angle. (A)** Average branching angle of auxin influx defective mutant *aux1-7* grown in 3%Glc alone or in combination with MeJA. **(B)** Distribution of branching angle of auxin influx defective mutant *lax3* grown in 3%Glc alone or in combination with different doses of 10 μM MeJA **(C)** Average branching angle of auxin efflux defective mutants *eir1-1* and *mdr1*.*1* grown in 3%Glc alone or in combination with 10 μM MeJA. **(D)** Distribution of branching angle of auxin efflux defective mutants *pin3-4, pin4-3* and *pin7-2* grown in 3%Glc alone or in combination with different doses of MeJA. **(E and F)** Average branching angle of auxin signalling mutants *tir1-1, axr1-3* and *axr2-1* grown in 3%Glc alone or in combination with MeJA. 5-day-old 1/2 MS grown Col-0 and mutant seedlings were transferred to treatment media and phenotypes were analysed at 12^th^ day. The data represents the average of 4 biological replicates consisting of 25 seedlings and error bars represent SE. Asterisks indicate a significant difference in the studied parameter (P <0.05, paired two-tailed student’s t-test; * control vs treatment and WT vs mutant).

**Figure S9. Role of PIN3 in JA mediated regulation of branching angle. (A)** Stage II LR of PIN3::PIN3-eGFP seedlings treated with 1/2 MS alone or in combination with 10 μM MeJA. **(B)** Quantification of fluorescence of stage II PIN3::PIN3-eGFP LRs treated with 1/2 MS alone or in combination with 10 μM MeJA. The graph represents average of 2 biological replicates with 10 roots imaged in each treatment.

**Figure S10. Role of light signalling in controlling JA-mediated branching angle. (A)** Phenotype of light and dark adapted Col-0 and *myc2myc3myc4* seedlings in different concentrations of Glc (0.5%, 3%) alone or in combination with MeJA (10 μM). **(B)** Distribution of branching angle in light and dark adapted Col-0 and *myc2myc3myc4* seedlings in different concentrations of Glc (0.5%, 3%) alone or in combination with MeJA (10 μM). The data represents the average of 6 biological replicates consisting of 25 seedlings and error bars represent SE. Asterisks indicate a significant difference in the studied parameter (P <0.05, paired two-tailed student’s t-test; * control vs treatment and WT vs mutant).

**Figure S11. Role of light on auxin distribution to regulate branching angle**. Distribution of branching angle in light and dark adapted Col-0 and *aux1-7* seedlings on ½ MS alone and in combination with MeJA (10μM). The data represents average of 4 biological replicates consisting of 25 seedlings and error bars represent SE. Asterisks indicate a significant difference in the studied parameter (*P* <0.05, Student’s t-test; *control vs treatment and WT vs mutant).

**Supplementary Table 1: Primers used in the study**.

## Notes

### Competing Interest Statement

The authors have declared no competing interest.

