## Supplementary data for "Jasmonic Acid coordinates with Light, Glucose and Auxin signalling in Regulating Branching Angle of *Arabidopsis* Lateral Roots"

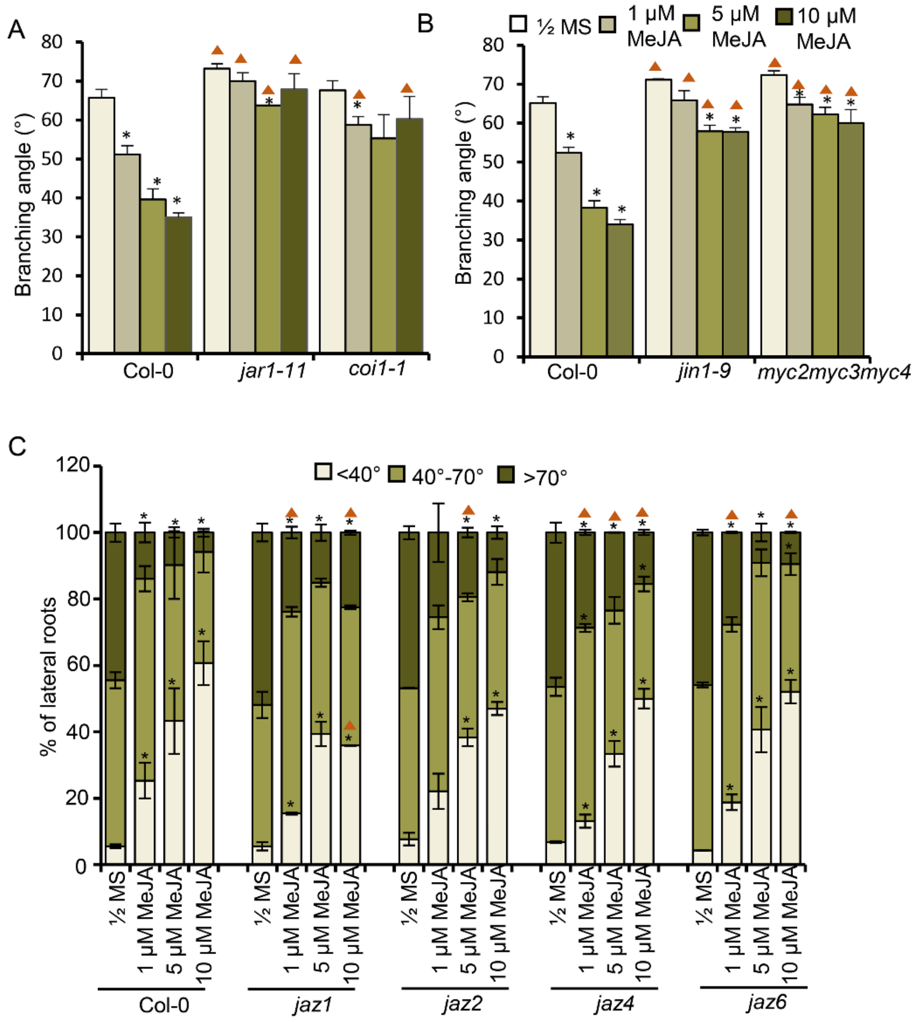


**Figure S1. MeJA regulation of branching angle of Arabidopsis roots. (A and B)** Average branching angle of 12-day-old Col-0 and JA signalling mutants grown in different doses of MeJA. The data represents the average of 4 biological replicates consisting of 25 seedlings and error bars represent SE. **(C)** Distribution of branching angle of Col-0 and different *jaz* seedlings grown in different doses of MeJA. The data represents the average of 3 biological replicates consisting of 25 seedlings and error bars represent SE. 5-day-old 1/2 MS grown Col-0 and mutant seedlings were transferred to treatment media and phenotypes were analysed at 12$^{th}$ day. Asterisks indicate a significant difference in the studied parameter (P <0.05, paired two-tailed student's t-test; * control vs treatment and ▲ WT vs mutant).

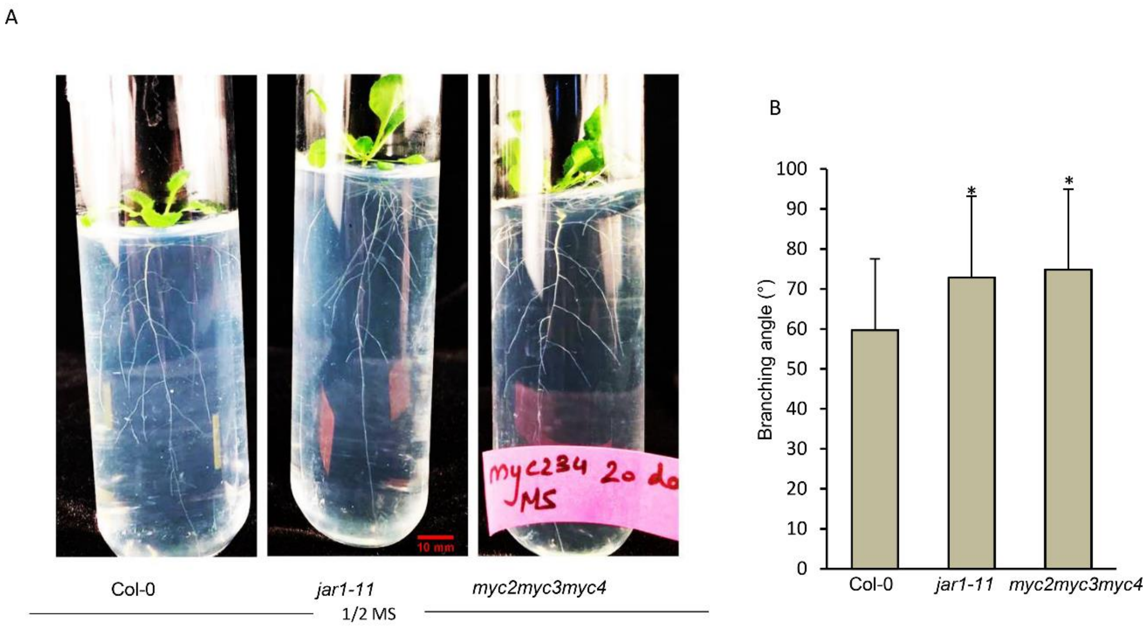


**Figure S2. MeJA regulation of branching angle of Arabidopsis roots. (A)** A representative image of 20 days after germination (DAG) of Col-0 and JA biosynthesis and signalling mutants *jar1-11* and *myc2myc3myc4* grown in 1/2 MS medium in cylindrical tubes. **(B)** Average branching angle of 20 DAG of Col-0, *jar1-11* and *myc2myc3myc4* grown in 1/2 MS medium in cylindrical tubes. The data represents images from 2 biological replicates, each having 5 plants (P <0.05, paired two-tailed student's t-test; * WT vs mutant).

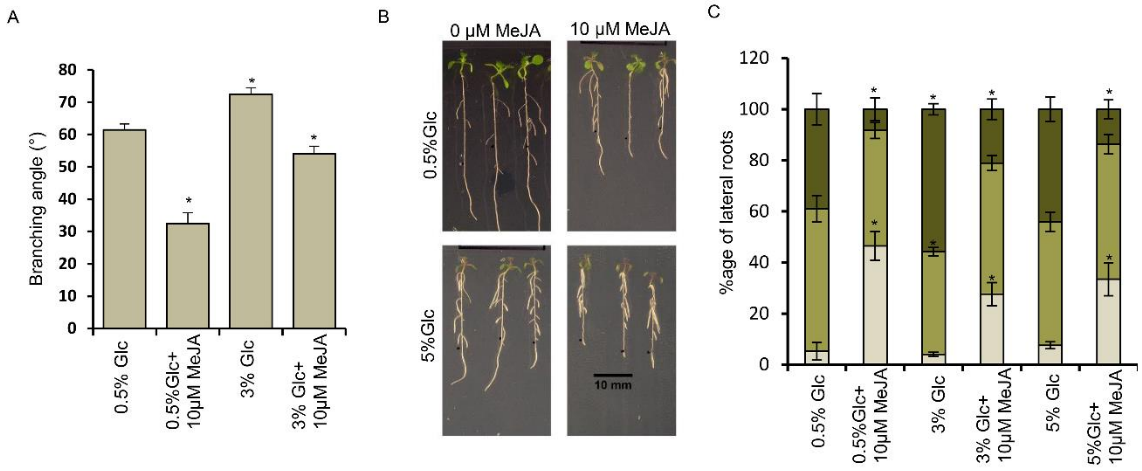


**Figure S3. MeJA-Glc regulation of branching angle of Arabidopsis roots.** **(A)** Average branching angle of 12-day-old Col-0 seedlings grown in different concentrations of Glc (0.5%, 3%) and in combination with 10 µM MeJA. The data represents the average of 7 biological replicates consisting of 25 seedlings and error bars represent SE (P<0.05, Student's t-test; * control (0.5%Glc) vs treatment. **(B)** Phenotype of Col-0 seedlings grown in different concentrations of Glc (0.5%; 28 mM and 5%Glc; 278 mM) and in combination with 10 µM MeJA. **(C)** Distribution of branching angle of seedlings of Col-0 grown in different different concentrations of Glc (0.5%, 3% and 5%) and in combination with MeJA. The data represents the average of 4 biological replicates consisting of 25 seedlings and error bars represent SE. Asterisks indicate a significant difference in the studied parameter (*P*<0.05,Student'st-test;*control (0.5%) vs treatment).

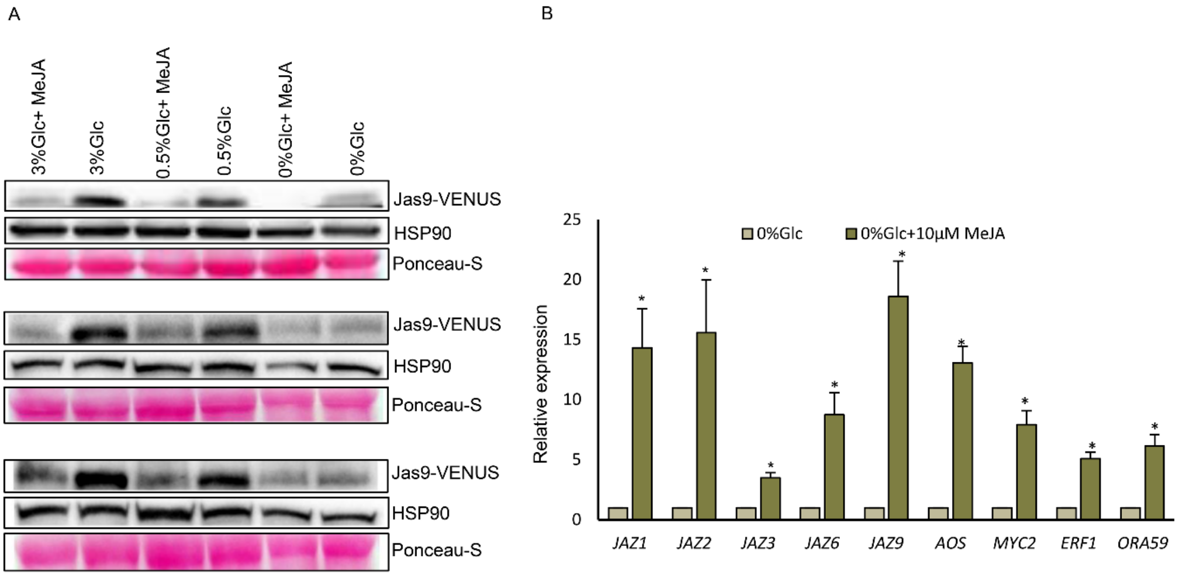


**Figure S4. MeJA-Glc regulation of branching angle of Arabidopsis roots. (A)** 3 individual biological replicates of Jas9-VENUS seedlings treated with increasing concentrations of Glc (0%Glc, 0.5%Glc, 3%Glc) along with 10 µM MeJA. **(B)** RT-qPCR showing the expression JA signalling genes (*JAZ1*, *JAZ2*, *JAZ3*, *JAZ6*, *JAZ9*, *AOS*, *MYC2*, *ERF1* and *ORA59*) in Col-0 seedlings grown in 0%Glc and 0%Glc+ 10 µM MeJA. 5-day-old 1/2 MS grown Col-0 seedlings were first starved for 24 hrs and transferred to liquid treatment media containing 10 µM MeJA and analysed after 3hrs. The data represents the average of 4 biological replicates consisting of 25 seedlings and error bars represent SE (P <0.05, paired two-tailed student's t-test; * control (0%Glc) vs treatment (0%Glc+MeJA).

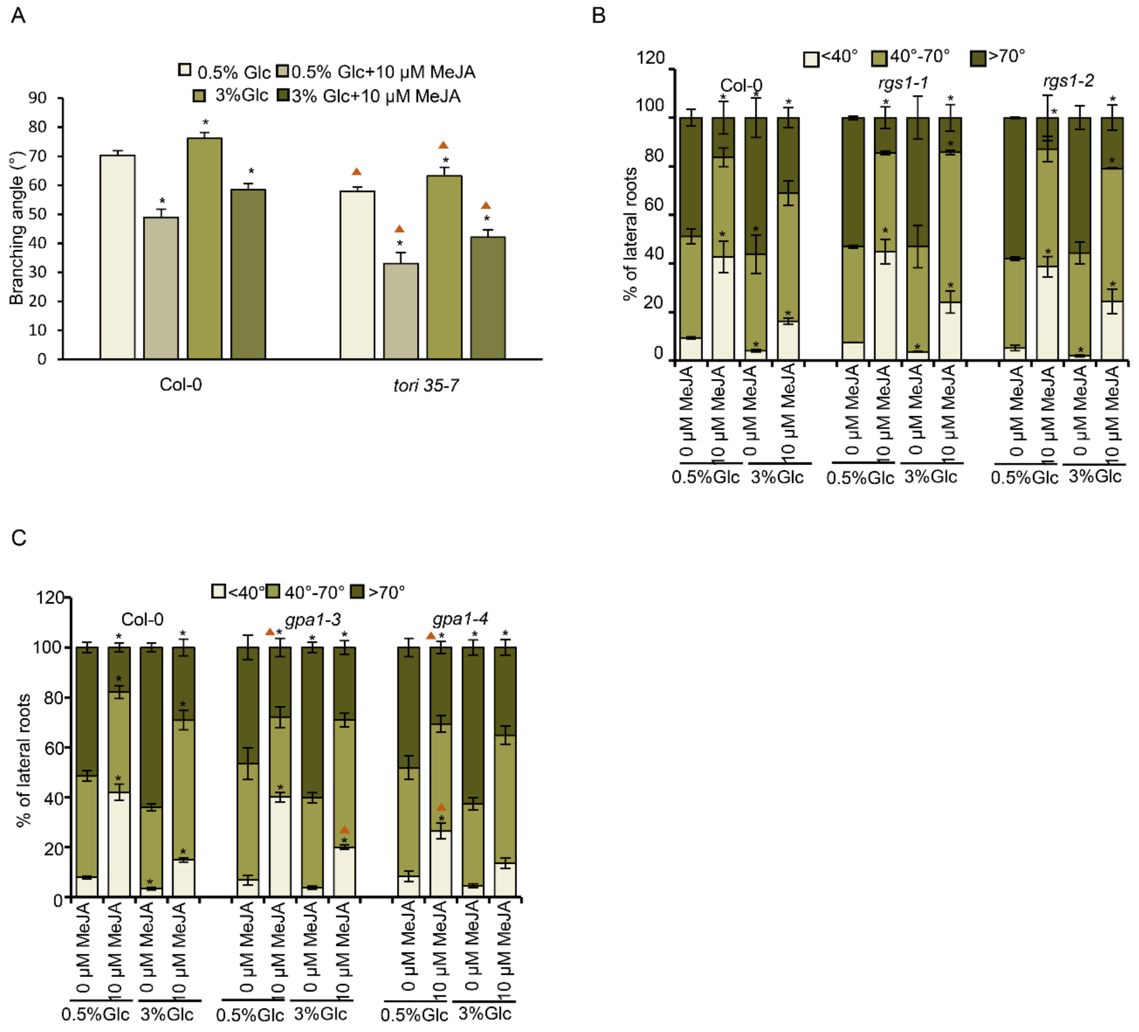


**Figure S5. Role of different components of Glc signalling in regulation of branching angle of Arabidopsis roots**. **(A)** Average branching angle of 12-day-old Col-0 and Glc-dependent TOR signaling mutant *tor35-7RNAi* grown in different concentrations of Glc (0.5%, 3%) and in combination with MeJA. The data represents the average of 4 biological replicates consisting of 25 seedlings and error bars represent SE. (*P* <0.05, Student's t-test; * control vs treatment ** 0.5%Glc vs 3%Glc+10MeJA and ▲wild-type vs mutant). **(B and C)** Distribution of branching angle in seedlings of Col-0 and RGS-dependent signalling mutants *rgs1-1*, *rgs1-2*, *gpa1-3* and *gpa1-4* grown in different concentrations of Glc in combination with MeJA. The data represents the average of 4 biological replicates consisting of 25 seedlings and error bars represent SE.

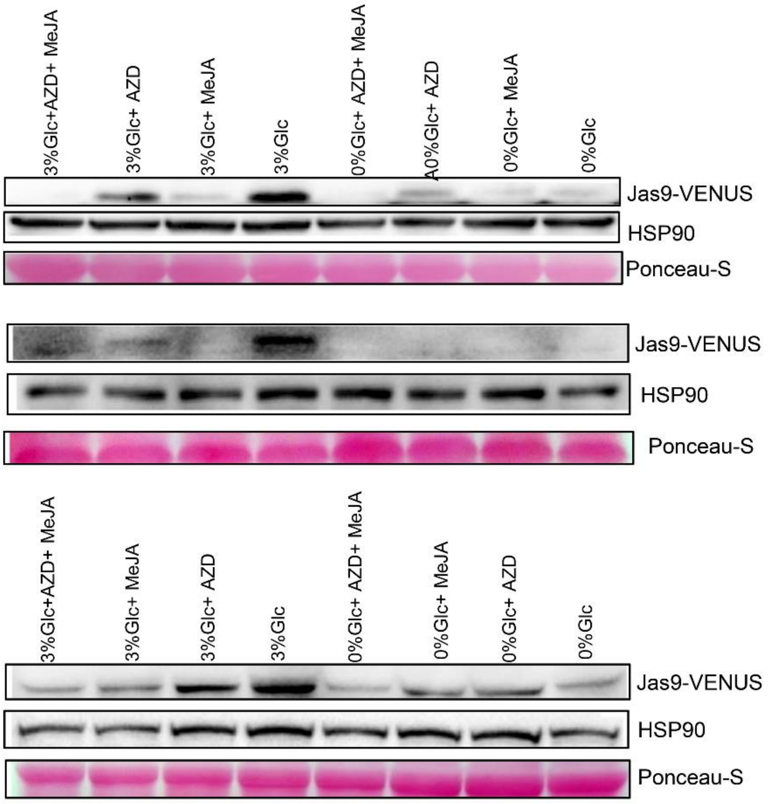


**Figure S6. Glc regulates branching angle of LRs via TOR-mediated signalling pathway.** 3 individual biological replicates of Jas9-VENUS seedlings treated without and with Glc (3%Glc) along with 10 µM MeJA, 10 µM AZD and a combination of 10 µM MeJA and 10 µM AZD (related to Figure 2J, K).

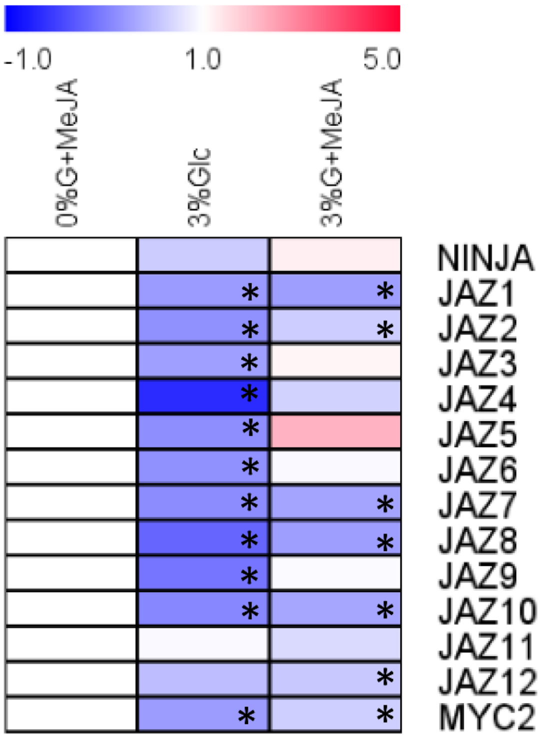


**Figure S7. Heat map of JA signalling genes upon treatment with 0%Glc+10µM MeJA, 3%Glc and 3%Glc+10µM MeJA.** The fold change value of genes (+/-1.5 fold) treated with 0%Glc+10µM MeJA was considered as 1. The fold change values of genes in 3%Glc and 3%Glc+ 10 µM MeJA was normalised (as % change) with those of 0%Glc+10µM MeJA. * indicates expression values of genes that are significantly altered. The data is the average of 2 biological replicates. Heat map was generated using MeV.

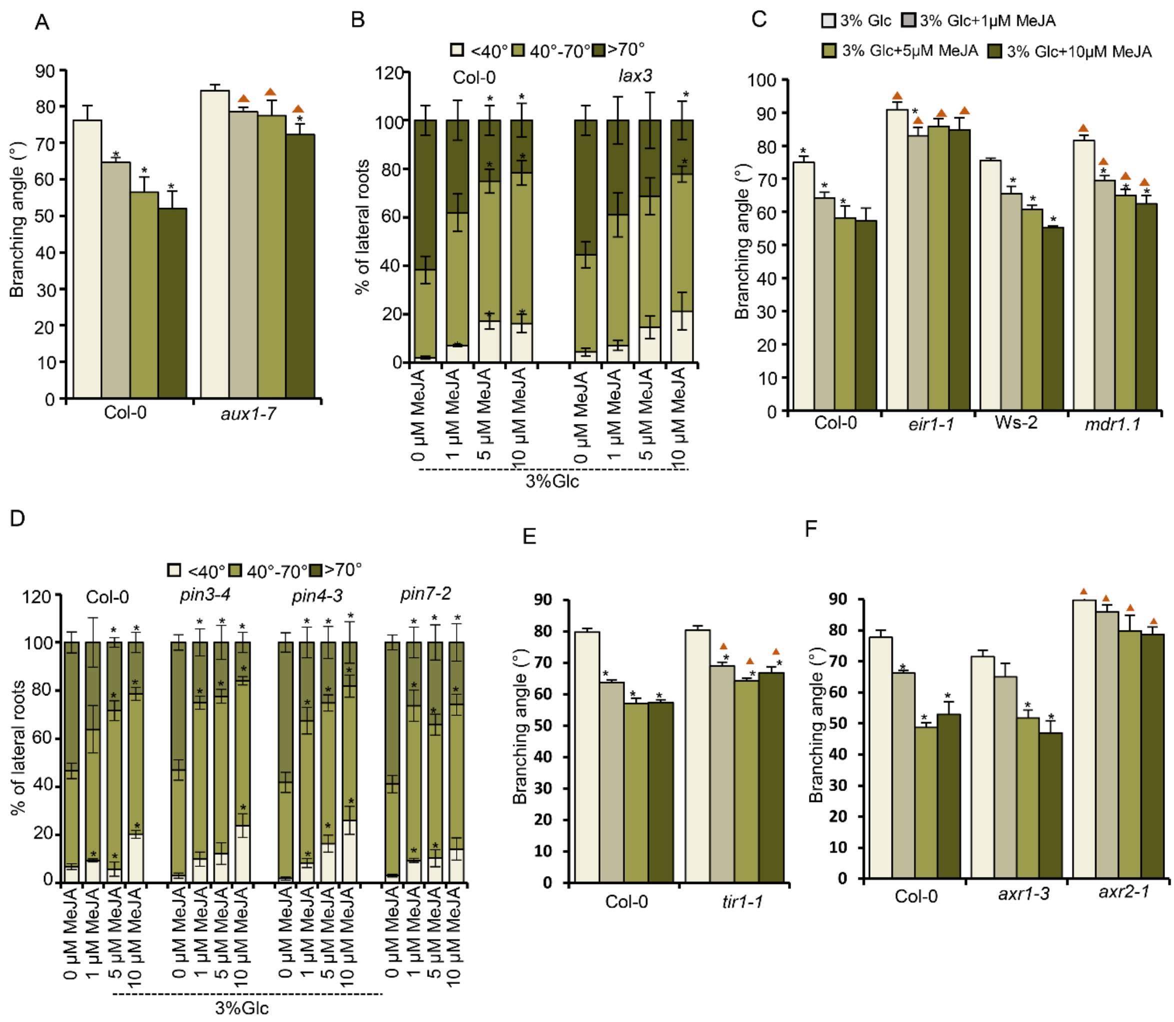


**Figure S8. Role of auxin machinery in controlling JA-mediated branching angle. (A)** Average branching angle of auxin influx defective mutant *aux1-7* grown in 3%Glc alone or in combination with MeJA. **(B)** Distribution of branching angle of auxin influx defective mutant *lax3* grown in 3%Glc alone or in combination with different doses of 10 µM MeJA **(C)** Average branching angle of auxin efflux defective mutants *eir1-1* and *mdr1.1* grown in 3%Glc alone or in combination with 10 µM MeJA. **(D)** Distribution of branching angle of auxin efflux defective mutants *pin3-4*, *pin4-3* and *pin7-2* grown in 3%Glc alone or in combination with different doses of MeJA. **(E and F)** Average branching angle of auxin signalling mutants *tir1- 1*, *axr1-3* and *axr2-1* grown in 3%Glc alone or in combination with MeJA. 5-day-old 1/2 MS grown Col-0 and mutant seedlings were transferred to treatment media and phenotypes were analysed at 12th day. The data represents the average of 4 biological replicates consisting of 25 seedlings and error bars represent SE. Asterisks indicate a significant difference in the studied parameter (P <0.05, paired two-tailed student's t-test; * control vs treatment and ▲WT vs mutant).

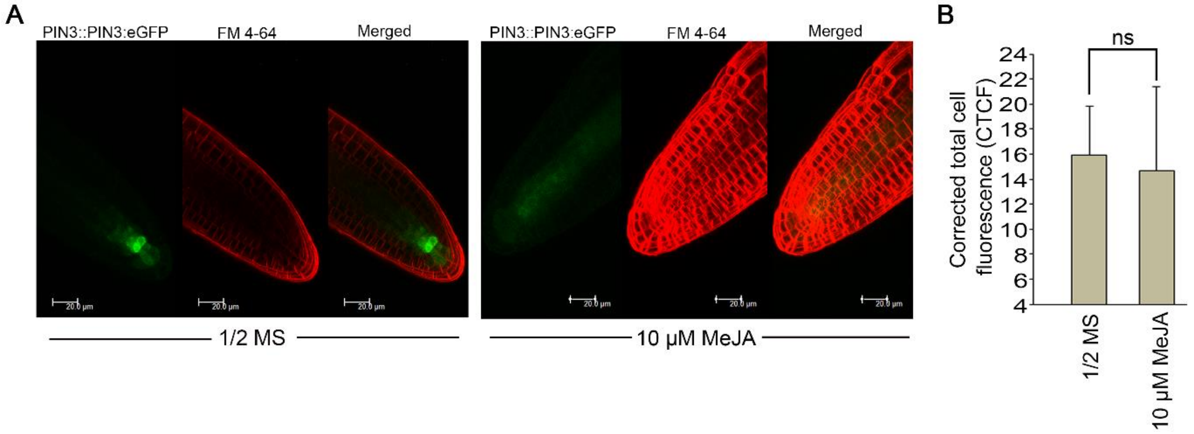


**Figure S9. Role of PIN3 in JA mediated regulation of branching angle. (A)** Stage II LR of PIN3::PIN3-eGFP seedlings treated with 1/2 MS alone or in combination with 10 µM MeJA. **(B)** Quantification of fluorescence of stage II PIN3::PIN3-eGFP LRs treated with 1/2 MS alone or in combination with 10 µM MeJA. The graph represents average of 2 biological replicates with 10 roots imaged in each treatment.

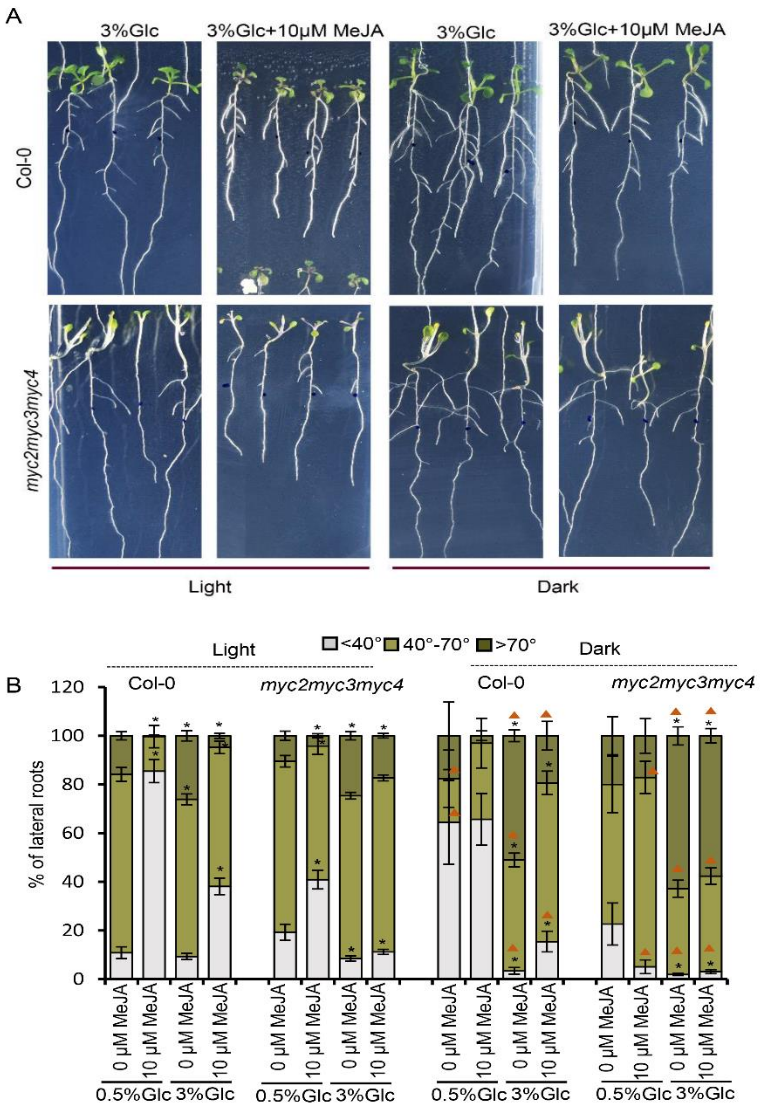


**Figure S10. Role of light signalling in controlling JA-mediated branching angle. (A)** Phenotype of light and dark adapted Col-0 and *myc2myc3myc4* seedlings in different concentrations of Glc (0.5%, 3%) alone or in combination with MeJA (10 µM). **(B)** Distribution of branching angle in light and dark adapted Col-0 and *myc2myc3myc4* seedlings in different concentrations of Glc (0.5%, 3%) alone or in combination with MeJA (10 µM). The data represents the average of 7 biological replicates consisting of 25 seedlings and error bars represent SE. Asterisks indicate a significant difference in the studied parameter ($P < 0.05$, paired two-tailed student's t-test; * control vs treatment and ▲ Light vs dark ).

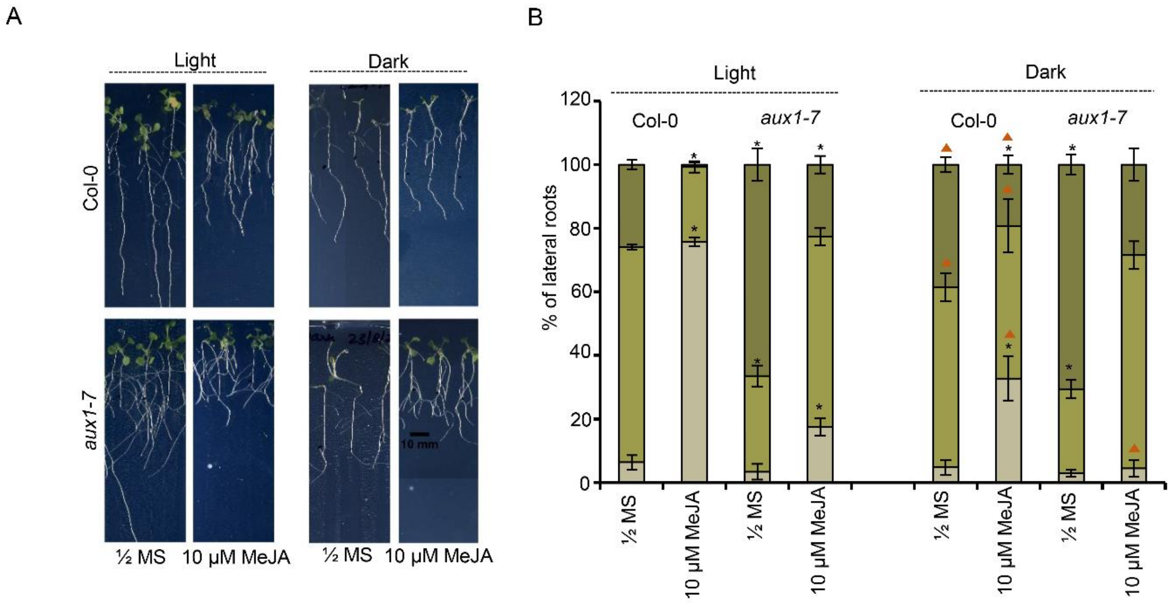


**Figure S11. Role of light on auxin distribution to regulate branching angle. (A)** Phenotype of light and dark adapted Col-0 and *aux1-7* seedlings in 1/2 MS and in combination with MeJA (10 µM). **(B)** Distribution of branching angle in light and dark adapted Col-0 and *aux1-7* seedlings on ½ MS alone and combination with MeJA (10µM). The data represents the average of 4 biological replicates consisting of 25 seedlings and error bars represent SE. Asterisks indicate a significant difference in the studied parameter ($P$ <0.05, Student's t-test; * control vs treatment and ▲ WT vs mutant).

| Oligo name | 5'-3' Sequence |
|---|---|
| MYC2_ChIP_CYP79B2 R1-F | CGTAGTCGTGTCATATAGCGATTAAC |
| MYC2_ChIP_CYP79B2 R1-R | GGATTGGTTTGTAATACTTTGCAAT |
| MYC2_ChIP_CYP79B2 R2-F | CATGGTGAAAAACATTTTGCTAGC |
| MYC2_ChIP_CYP79B2 R2-R | CCCAAATGTTTTCGCCTTCT |
| MYC2_ChIP_LZY2 R1-F | GGCCATAGAAGAATGTTGGTGGA |
| MYC2_ChIP_LZY2 R1-R | CAAGTTCCAAACTTTGACTACTTAAAAGAGA |
| MYC2_ChIP_LZY2 R2-F | GATTCATCAATAACTCGAGTTGAGAAAT |
| MYC2_ChIP_LZY2 R2-R | GAAAACAAAAGCCAGAAGAAAACACTT |
| MYC2_ChIP_LZY4 R1-F | GCGCTTCTAAGTTATAACTATACACTGATT |
| MYC2_ChIP_LZY4 R1-R | GAGGCTTCCACACATTTTGTTG |
| ATXR6_ChIP FP | CCGAACCGAACAACCAAAATATATG |
| ATXR6_ChIP RP | CCAGAGAAAGAGAGAGAGTGAGAGATT |
| ORA59_ChIP FP | GTACGTCATACACTCAACCTG |
| ORA59_ChIP RP | CAATTAGGCTGCCTCCGAATA |
| LOX3 RT-qPCR-F | ACGCTGATCCTGACCGTAGAA |
| LOX3 RT-qPCR-R | GCTCAGAACTCGGAACCAACA |
| AOS RT-qPCR-F | CAACCCCTTTTCCGATTTCTC |
| AOS RT-qPCR-R | CGGAATTTCAACGGCTTTGA |
| JAZ1 RT-qPCR-F | GCCTGTCTAAATCCCCTTGCT |
| JAZ1 RT-qPCR-R | AGAGTATTTGATAGTATGGTTCGTCAACA |
| JAZ2 RT-qPCR-F | GCTTGGCTCAGTTCACGGTAA |
| JAZ2 RT-qPCR-R | GGTCTCTCAAGTTGTGCCTTCTG |
| JAZ3 RT-qPCR-F | TCCACATCCATATTTGCATTTCA |
| JAZ3 RT-qPCR-F | CGAGTGAGGAAAAATTAACTGAGACA |
| JAZ6 RT-qPCR-F | CAACAACACATGAGCAAAAAGCT |
| JAZ6 RT-qPCR-R | TTTACACGTACAAAAAATCGAGGATT |
| JAZ9 RT-qPCR-F | ACGAGCAGGAGAAGACGTTAGG |
| JAZ9 RT-qPCR-R | CTACAATAAACAGACCAAAGCATTACAA |
| MYC2 RT-qPCR-F | CTCAAGCACAATAACCGAAAACC |
| MYC2 RT-qPCR-R | TTCGGATTCTGGGTCTGAGAA |
| CYP79B2 RT-qPCR-F | GGCTCCGGCGCTAGGA |
| CYP79B2 RT-qPCR-R | TTGAAGAAGTCTCGCGAGCAT |
| LAZY2 RT-qPCR-F | AGAGGAATTTTTTTGGTGCATCTG |
| LAZY2 RT-qPCR-R | CAGATCAAATCATGCAAAAAAAGAAG |
| LAZY4 RT-qPCR-F | TTTTTCCACCTCTTTCTTTGTCTTC |
| LAZY4 RT-qPCR-R | TCCACCCGAAAAACTTCATGT |
| ERF1 RT- qPCR -F | TCTTTGAGGATTTGGGAGAACAG |
| ERF1 RT- qPCR -R | CCAAGTCCCACTATTTTCAGAAGAC |
| ORA59 RT- qPCR -F | TGTTCTTGATAATCTCTGCTTCTACAATT |
| ORA59 RT- qPCR -R | GCCATCACATCTCTTCCAAAACT |

**Supplementary Table 1: Primers used in the study.**
